# Impact of Chemical Dynamics of Commercial PURE Systems on Malachite Green Aptamer Fluorescence

**DOI:** 10.1101/2024.03.15.585317

**Authors:** Zoila Jurado, Richard M. Murray

## Abstract

The malachite green aptamer (MGapt) is known for its utility in RNA measurement *in vivo* and lysate-based cell-free protein systems. However, MGapt fluorescence dynamics do not accurately reflect mRNA concentration. Our study finds that MGapt fluorescence is unstable in commercial PURE systems. We discovered that the chemical composition of the cell-free reaction strongly influences MGapt fluorescence, which leads to inaccurate RNA calculations. Specific to the commercial system, we posit that MGapt fluorescence is significantly affected by the system’s chemical properties, governed notably by the presence of dithiothreitol (DTT). We propose a model that, on average, accurately predicts MGapt measurement within a 10% margin, leveraging DTT concentration as a critical factor. This model sheds light on the complex dynamics of MGapt in cell-free systems and underscores the importance of considering environmental factors in RNA measurements using aptamers.

## Introduction

Cell-free protein synthesis (CFPS) systems can be broadly categorized into two types. The first and most commonly used CFPS system is based on cell lysate. The cell lysate-based system uses cellular machinery harvested from the cell [1]. The second category contains all the necessary transcription and translation proteins for *E. coli*, each cultured and purified individually and combined at known concentrations, known as PURE — Protein synthesis Using purified Recombinant Elements [2]. A variant of PURE is OnePot PURE, where all 36 proteins are co-cultured and purified together [3].

Using PURE facilitates the modeling of the CPFS system, allowing for the development of a complete model and techniques to seamlessly integrate it into the “design-build-test” pipeline for genetic circuit construction or synthetic cell assembly. Models in recent years have modeled the translation of peptides for PURE using chemical reaction networks [4, 5], and our previous work has added to these models by expanding the user peptide and the addition of transcription [6]. Validation of all models requires accurate transcription and translation monitoring. While translation can be measured using fluorescent proteins, transcription is more challenging, leading to the reliance on RNA aptamers, such as malachite green aptamer. Initially used as a means to control gene expression in *S. cerevisiae* [7], MGapt has been used as a means to study RNA production, RNA dynamics, and investigating trade-offs between transcription and translation in cell-free protein systems with protein expression of deGFP [8, 9, 10].

Studies using MGapt to measure RNA production in CFPS have been used in lysates [9, 11]. Our use of MGapt in PURExpress reveals surprising dynamics, which may explain why the MGapt measurements appear absent from PURE CFPS. Generally, the exceptional specificity of aptamers allows the discrimination between closely related isoforms or different conformational states of the same target molecule [12, 13]. However, MGapt has been known to bind to other triphenylmethane dyes such as crystal violet (CV), tetramethylrhodamine (TMR), and Pyronin Y (PY) [14, 15]. Furthermore, it has been found that free MGapt reduces the amount of RNA folded in the correct binding conformation, and metal ions, though not required for high-affinity bind [16], stabilize the complexes with non-native ligands. In contrast, the complex with the original selection target is stable at low salt and without divalent metal ions [17]. The destabilization of MGapt through the use of organic solvent, the addition of Mg^2+^, DTT, and other ions has been described in the past [18, 19, 20, 21, 15, 22].

In this paper, we demonstrate how the chemical composition of commercial PURE may destabilize MGapt, leading to different aptamer states corresponding to different fluorescence levels. We will first describe the observed MGapt expression in commercially available PURE and inconsistencies between what we could predict and what was observed, such as saturation time and dynamics of MGapt when no transcription is involved. We then discuss the potential effects of DTT in the system and propose a model that uses DTT as a driving force. Finally, we provide experimental validation of the measured MGapt model of the PURE system accounting for DTT’s impact on the state of MGapt, allowing for accurate RNA calculations.

## Results and Discussion

### Expression and fluorescence of MGapt in different PURE systems

We initially evaluated the consistency of MGapt dynamics across two distinct PURE systems to ascertain whether this phenomenon is inherent to PURE. In commercial PURE systems, the protocol recommends incubating the reaction for 2-4 hours [23, 24], during which protein production saturates. RNA production would also be expected to saturate around the 2-hour mark to be consistent with the reaction lifespan. However, MGapt fluorescence does not saturate at 2 hours. In comparison in PURExpress and OnePot PURE, both expressing MGapt from plasmid DNA with construct pT7-MGapt-tT7 (light blue) and pT7-MGapt-UTR1-deGFP-tT7 (dark blue), MGapt fluorescence dynamics were striking, shown in Figure 1.

**Figure 1:**
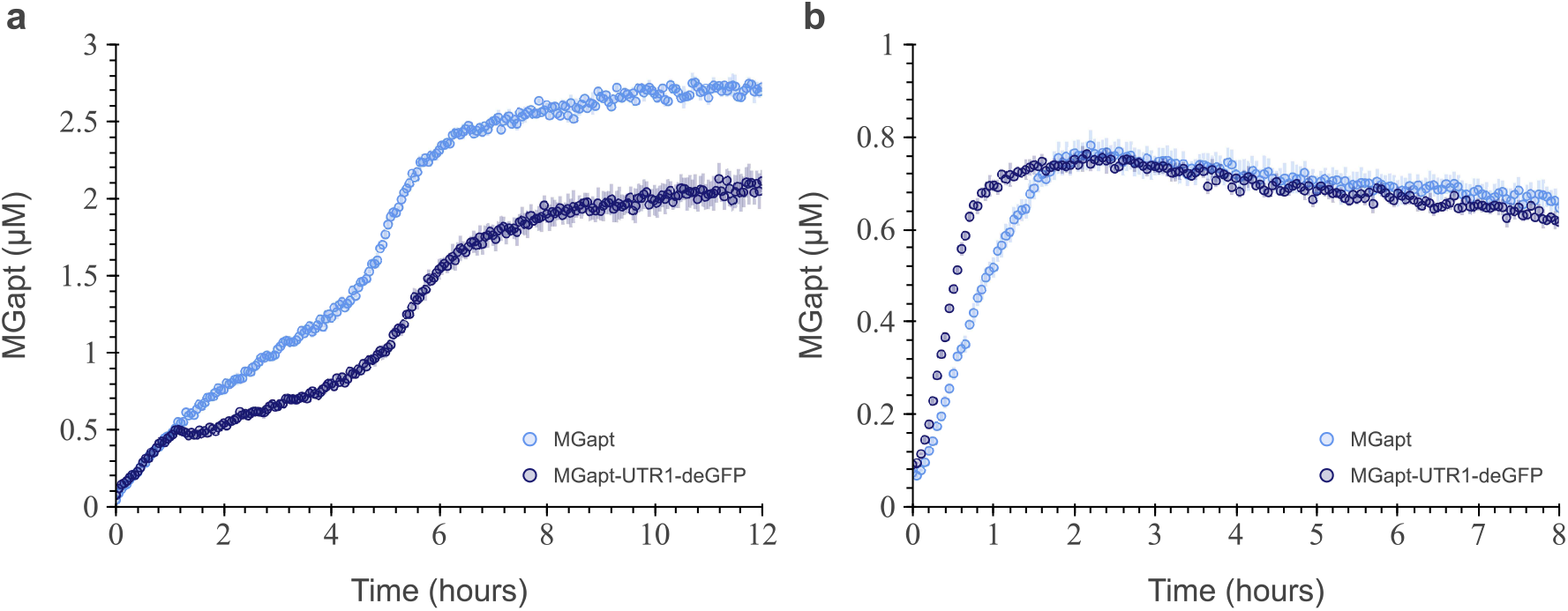
MGapt concentration in PURExpress and OnePot CFPS system. The MGapt concentration over time for transcriptions of pT7-MGapt-tT7 (light blue) and pT7-MGapt-UTR1-deGFP-tT7 (dark blue) at 5 nm, in (**a**) PURExpress and (**b**) OnePot, made in lab. Experimental data consisted of three replicates (circles and respective error bars).

Experimental results indicate that the two PURE systems had different MGapt dynamics, irrespective of the total MGapt production. While each system exhibited a monotonic upward trend of MGapt fluorescence, the PURExpress reaction (Figure 1a) showed that MGapt fluorescence saturated at around six hours. On the other hand, the OnePot PURE reaction (Figure 1b) showed that MGapt fluorescence saturated at around two hours. The MGapt saturation could indicate that MGapt production continues after two hours in PURExpress, effectively out-living OnePot PURE’s RNA production. It didn’t appear logical for the transcription process to persist beyond the translation. Furthermore, if MGapt fluorescence is considered an accurate reflection of RNA production, it would also suggest that the RNA production rate changes around four hours. The significant disparity in MGapt production dynamics suggested MGapt fluorescence might increase due to chemical differences between PURExpress and OnePot PURE.

### Effects of buffering conditions MGapt fluorescence

Upon investigating the difference, we first the two systems chemical compositions [2, 25]; outlined in Table S1. We discovered numerous salts in one system but not the other, which we concluded should not significantly impact MGapt fluorescence. The one major difference in the two systems was the reducing agent used, tris(2-carboxyethyl)phosphine (TCEP) versus dithiothreitol (DTT). PURExpress uses 1 mM of DTT [26] while OnePot PURE uses 1 mM of TCEP [3]. DTT and TCEP both reduce disulfide bonds, but TCEP has the advantages of being significantly more stable in the absence of a metal chelator and less inhibitive in labeling with maleimide [27]. Additionally, compared to DTT, TCEP is more effective, non-volatile, and does not readily oxidize above pH 7.5 [28]. It is important to note that the PURE reaction occurs above pH 7.5, Figure S1.

MGapt fluorescent was measured under four different chemical conditions to measure the effect of the reducing agent on MGapt fluorescence in separate *in vitro* reactions. Three different reducing agents were tested: TCEP, glutathione (GSH), and DTT), and the waste product pyrophosphate (PPi). The experiment consisted of three replicates of 10 µL reactions with the respective additives (TCEP, GSH, DTT, and PPi) at experimentally relevant concentrations and purified RNA of MGapt-UTR1-deGFP at 0.51 µm. MGapt fluorescence was read in BioTek plate reader (610/650) for six hours at 37 °C and calibrated to µm using calibration curve Figure S2. Figure 2 summarizes the average MGapt concentration measured over the six-hour window; full-time course measurements are depicted in Figure S3.

**Figure 2:**
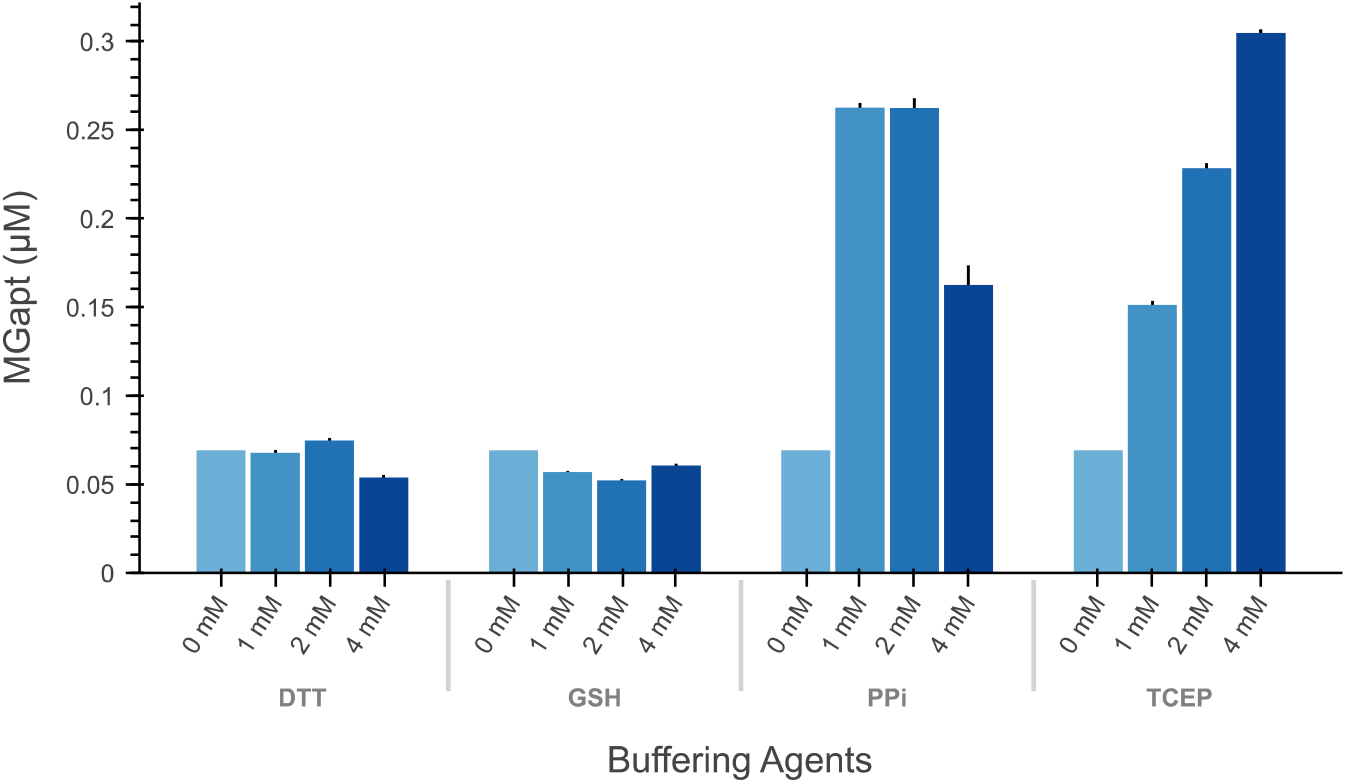
Measured MGapt concentration under different buffer conditions. Average measured MGapt concentration of purified RNA coding for MGapt-UTR1-deGFP in different reducing agents and waste chemicals over six hours. Each section of the bar plot is titled with the respective buffer used, and the increasing chemical concentrations are designated with increasing intensity of blue. The negative control was subtracted from the test conditions, and standard deviations of triplicates are represented by black bars.

Only TCEP and PPi samples showed concentration effects on the total MGapt fluorescence. Higher concentrations of TCEP and PPi resulted in higher MGapt fluorescence, except at 4 mM PPi, where MGapt fluorescence decreased. The results from Figure 2 did not provide conclusive evidence that the reducing agent, DTT, directly affected MGapt fluorescence nor that other reducing agents (TCEP or GSH) would be a better alternative.

### The fluorescence of MGapt is affected by the chemical properties of the PURE system

While Figure 2 implies that the fluctuating concentration of PPi might lead to increased MGapt fluorescence, it’s important to note that waste production would be directly correlated with system expression. Therefore, we wouldn’t anticipate a greater or quicker production of PPi in the PURExpress than in OnePot PURE. To avoid any possible interference of transcription products with MGapt, purified RNA of MGapt-UTR1-deGFP was added, removing the largest PPi production source, RNA strand elongation. Additionally, to isolate the effect of the PURE reaction on MGapt fluorescence, the translation reactions were minimized by using RNA containing only the MGapt.

Upon first observation, in both cases with 0.49 µm of MGapt-deGFP-RNA (crosses) or 0.53 µm of MGapt-RNA (circles), the measured MGapt concentration is not constant and that the RNA only containing MGapt measures lower than the RNA of MGapt-UTR1-deGFP. However, when we normalize each result by its respective maximum, Figure 3a, the concentration of MGapt appears to increase gradually over time despite the absence of transcription. Interestingly, the environmental effect seemed similar for both the TL-null and TL-only conditions, indicating that the additional ribosome binding site and deGFP sequences could not solely affect MGapt fluorescence.

**Figure 3:**
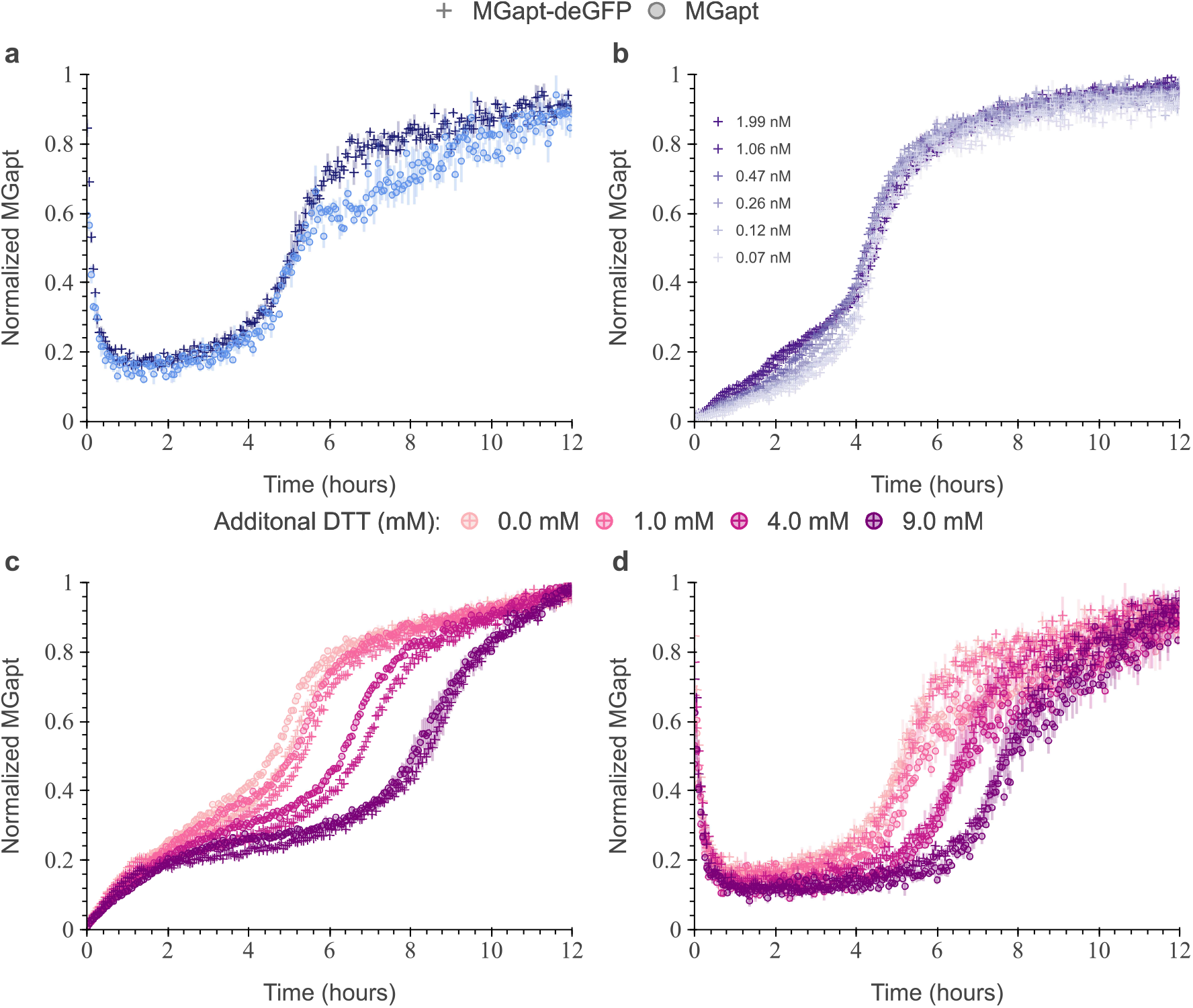
Normalized measured MGapt concentration dynamics in PURExpress. PURExpress reactions of 10 µL with three technical replicates containing 10 µm of malachite green dye and 8 units of RNAse inhibitor. (**a**) Starting from purified RNA of MGapt at 0.53 µm (light blue crosses) and MGapt-UTR1-deGFP at 0.49 µm (dark blue circles) after data is normalized. (**b**) Expressing plasmid pT7-MGapt-UTR1-deGFP-tT7 at various DNA concentrations after data is normalized. (**c**) Expressing of plasmid pT7-MGapt-UTR1-deGFP-tT7 at 4.93 nm and pT7-MGapt-tT7 at 5.03 nm with increasing DTT concentration after data is normalized. (**d**) Starting from purified RNA of MGapt-UTR1-deGFP at 0.49 µm and MGapt at 0.53 µm with increasing DTT concentration after data is normalized. Non-normalized measured MGapt concentration data are shown in Figure S4.

To further demonstrate that the measured MGapt concentration over time was not dependent on the production of RNA or protein, additional PURE reactions were performed at different DNA concentrations. The experiment consisted of three replicates of 10 µL with DNA plasmid pT7-MGapt-UTR1-deGFP-tT7 at six different DNA concentrations dispensed using an Echo 525 Acoustic Liquid Handler and the final DNA concentration was calculated on the rounded amount of DNA that was dispensed. To account for total RNA production, the measured MGapt concentrations were normalized, Figure 3b. The initial RNA production rate varied based on DNA concentrations, but all normalized MGapt concentration measurements converged. Notably, all concavity transitions occurred around the 4-hour mark regardless of RNA production rate, indicating that MGapt dynamics are unaffected by DNA concentration. The increase in MGapt concentration over time further supports that MGapt fluorescence is affected by the chemical environment of the commercial PURE system, independent of transcription and translation.

The observed inflection point around 4 hours, in Figures 3a-b, suggests the presence of system repression by specific chemical substrates that degrade over time. As illustrated in the model for PURE translation [4], there are 483 reactions with a nonzero rate, and 278 reactions occur without the presence of DNA: tRNA charging, NTP degradation, and energy recycling. These reactions also occur in OnePot PURE, indicating that this MGapt response wouldn’t be exclusive to one system. This leads back to the distinctions between OnePot PURE and commercial PURE, particularly their selection of reducing agents—DTT versus TCEP. While the reducing agent may not directly impact MGapt fluorescence, it could potentially influence other chemical reactions that affect MGapt fluorescence. To test the effects of DTT on measured MGapt concentration, DTT was added to PURExpress expressing DNA plasmids of pT7-MGapt-UTR1-deGFP-tT7 and pT7-MGapt-tT7 at 5 nm. DTT was added to reach a final added DTT concentration of 1 mM, 4 mM, and 9 mM. Following data normalization, Figure 3c, results showed that increasing DTT concentrations shifted the infection point of measured MGapt to the right. Additionally, we observed that the measured MGapt concentrations for both plasmids overlap, indicating that MGapt effects are primarily due to the addition of DTT.

Finally, the previous experiment was repeated with purified RNA to underscore the concept that DTT has a direct impact and can effectively represent other auxiliary reactions and chemicals affecting MGapt. In this iteration, purified RNA of MGapt-UTR1-deGFP and MGapt at approximately 0.5 µm concentration was combined with varying concentrations of DTT. In the normalized data, Figure 3d, data collapse based on added DTT concentration. Consistent with previous findings, as the concentration of DTT increased, the addition of DTT seemed to inhibit MGapt fluoresces. Based on these experimental results, we conclude that DTT acts as a direct or indirect suppressor of MGapt fluorescence, which degrades over time due to its volatile nature. This signifies that using MGapt measurements as an indicator for RNA is inaccurate as it does not account for DTT’s effect.

### MGapt transitions different aptamer states with varying levels of fluorescence

We assert that MGapt in the commercial PURE reaction goes through different equilibrium and intermediates states, as MGapt has been previously reported [22, 15, 19]. The concentration of DTT plays a crucial role in meditating, promoting, and/or facilitating the instability of MGapt and the conversion between the two chemical states, as demonstrated in Figure 3c-d. Using DTT as a proxy for the full chemical reactions responsible for altering MGapt states and thus fluorescence in the commercial PURE reaction, we propose the model depicting MGapt inhibition by DTT, as illustrated in Figure 4.

**Figure 4:**
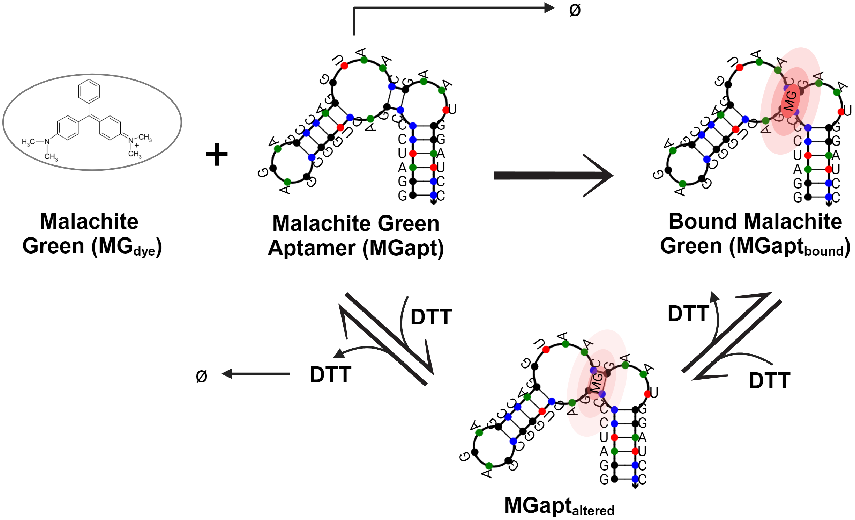
Inhibition of malachite green aptamer by DTT. Illustration of proposed interaction of DTT on bound fluorescent malachite green aptamer (MGapt). Adapted from chemical structure Stead *et al*. [19] and malachite green aptamer NuPack predicted secondary structure [29]. Created with BioRender.com

To model the detailed step-by-step inhibition and conversion of MG_dye_ bounded to MGapt (MGapt_bound_) to a less fluorescent MGapt_altered_, we built a chemical reaction network (CRN) in a CRN compiler tool called BioCRNpyler [30]. We utilize the functionalities provided by BioCRNpyler model the reactions shown in Figure 4:

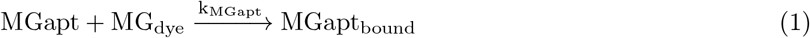

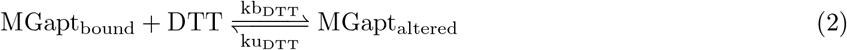

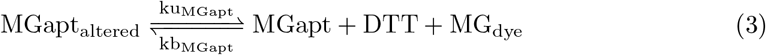

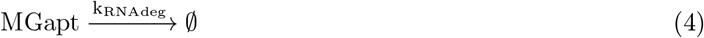

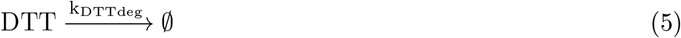

Beginning with transcribed and folded malachite green aptamer state (MGapt), malachite green oxalate (MG_dye_) binds to MGapt to form fully fluorescent malachite green aptamer (MGapt_bound_). Concentrations of DTT induce instability in MGapt_bound_ through a reversible process, leading to an intermediate state with reduced fluorescence (MGapt_altered_). The MGapt_altered_ state can be further destabilized, causing dissociation into its three components: MG_dye_, MGapt, and DTT. Finally, unbound states such as MGapt and DTT are susceptible to degradation. The introduction of the altered state, MGapt_altered_, which exhibits lower fluorescence compared to MGapt_bound_, implies that the total MGapt fluorescence is a linear combination of both states, MGapt_measured_ = MGapt_bound_ + c_1_MGapt_altered_. The MGapt_measured_ represents the measured fluorescence detected by the BioTek plate reader or any other fluorescent reader.

### Analysis and identification of parameters in the proposed model

To analyze the model of the effects of DTT on MGapt measured, we measured the MGapt fluorescence in a 10 µL reaction done in triplicate using PURExpress. The reaction contained 0.22 µm of RNA of MGapt-UTR1-deGFP. To determine the coefficients of the linear combination, we utilized the lowest concentration measurement to indicate when 100% of the aptamer was in the MGapt_altered_ state. The coefficient c_1_ was then determined based on the percentage of MGapt_measured_ and total added RNA (MGapt_total_), resulting in the final linear system:

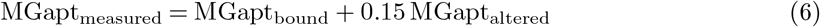

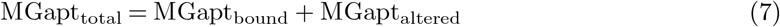

To parameterize the CRN model constructed using BioCRNpyler, we initialized the parameters by hand-tuning the reactions for bounded conditions by steps. Due to the time required to prepare the reactions before measuring on BioTek, it was necessary to determine the initial conditions at the time of reading. Employing the system of equations, we computed the initial conditions based on the MGapt_measured,t=0_ reading. Our calculations revealed that approximately 45% of the system was in the MGapt_bound_ state, while another 55% was in the MGapt_altered_ state. The specific values of the initial conditions utilized are outlined in Table S2. For accurate predictions of the measured MGapt concentration in PURE from the BioTek readings, we conducted model training using experimental data from the measured MGapt concentration of 0.22 µm RNA with the MGapt-UTR1-deGFP construct. Training this model using parameter identification was challenging due to the combined fluorescence of two distinct states in the measured data. With the ability to train only on one species, the effective concentration of MGapt_bound_ was calculated using the set of linear equations. Ultimately, in addressing the inherent noise in the experimental data, we employ Bayesian inference to derive a distribution of potential parameter values based on the experimental data. To accomplish both tasks of evaluating identifiability and determining posterior parameter distributions, we utilize a biological data analysis pipeline [31] implemented with the Python package Bioscrape [32].

Parameters for identification were determined by conducting a local sensitivity analysis of all species in the model across all parameters and time. The sensitivity analysis heatmap for DTT’s interaction with RNA of MGapt-UTR1-deGFP model is depicted in Figure 5a. We selected parameters with the highest sensitivity regarding MGapt output fluorescence. Among the seven reaction rates in equations (1)-(5), we identify k_MGapt_, kb_DTT_, ku_MGapt_, kb_MGapt_, and k_DTTdeg_ as the most sensitive parameters.

**Figure 5:**
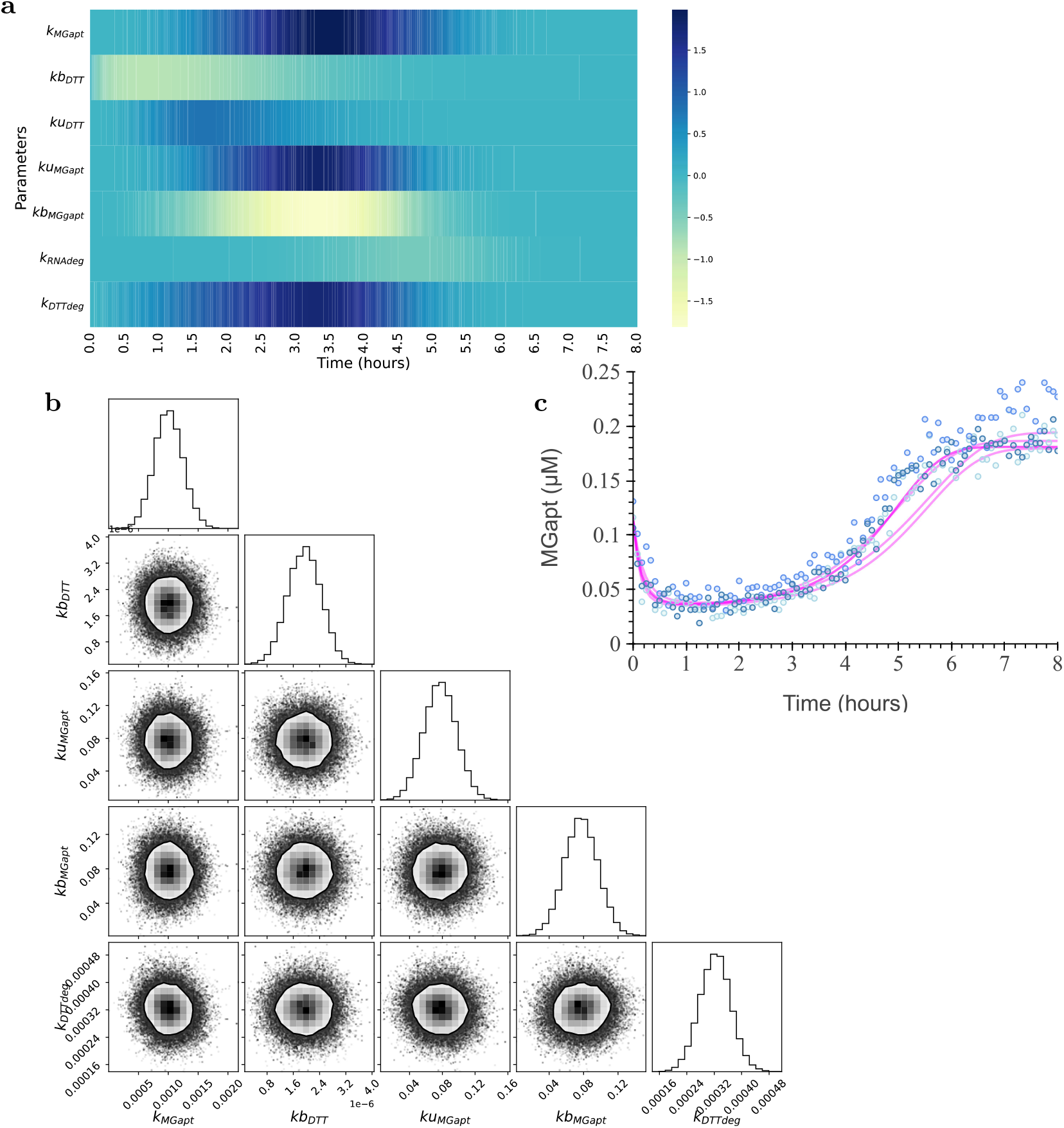
Modeling, analysis, and parameter inferencing of DTT effects on MGapt model. (**a**) The sensitivity of the MGapt to the CRN model parameters for the initial four hours. (**b**) The posterior distributions of parameters obtained after running Bayesian inference on k_MGapt_, kb_DTT_, ku_MGapt_, kb_MGapt_, and k_DTTdeg_. The corner plot depicts the covariance of the two parameters, with the contour showing the 75% probability region for the parameter values. (**c**) With parameter values for k_MGapt_, kb_DTT_, ku_MGapt_, kb_MGapt_, and k_DTTdeg_ sampled from the posterior distributions, the five model simulations (magenta lines) are shown alongside the experimental data for three biological replicates (scattered blue points).

Subsequently, we utilize Bayesian inference tools in Bioscrape to ascertain the posterior parameter distributions for these five parameters. The corner plot in Figure 5b illustrates the posterior parameter distributions and their covariance, providing a sampling distribution for predicting output using the fitted model. Model simulations using parameter values drawn from the posterior, alongside experimental data, are presented in Figure 5c. The final parameter values utilized are detailed in Table S4.

### Validation and assessment of model capturing DTT’s impact on MGapt fluorescence

To validate the model of the effects of DTT on MGapt measured, we conducted simultaneous tests using multiple RNA concentrations. The RNA of MGapt-UTR1-deGFP at final concentrations of 0.41 µm, 0.86 µm, 1.26 µm and 1.67 µm were read for 12 hours at 37 °C in a BioTek reader at 610/650 (ex/em). The initial concentrations of MGapt and MGapt_altered_ at *t* = 0 were determined as previously outlined and are listed in Table S3. Finally, utilizing equation (6), the MGapt_measured_ was computed and superimposed onto the experimental results using their respective colors shown in Figure 6.

**Figure 6:**
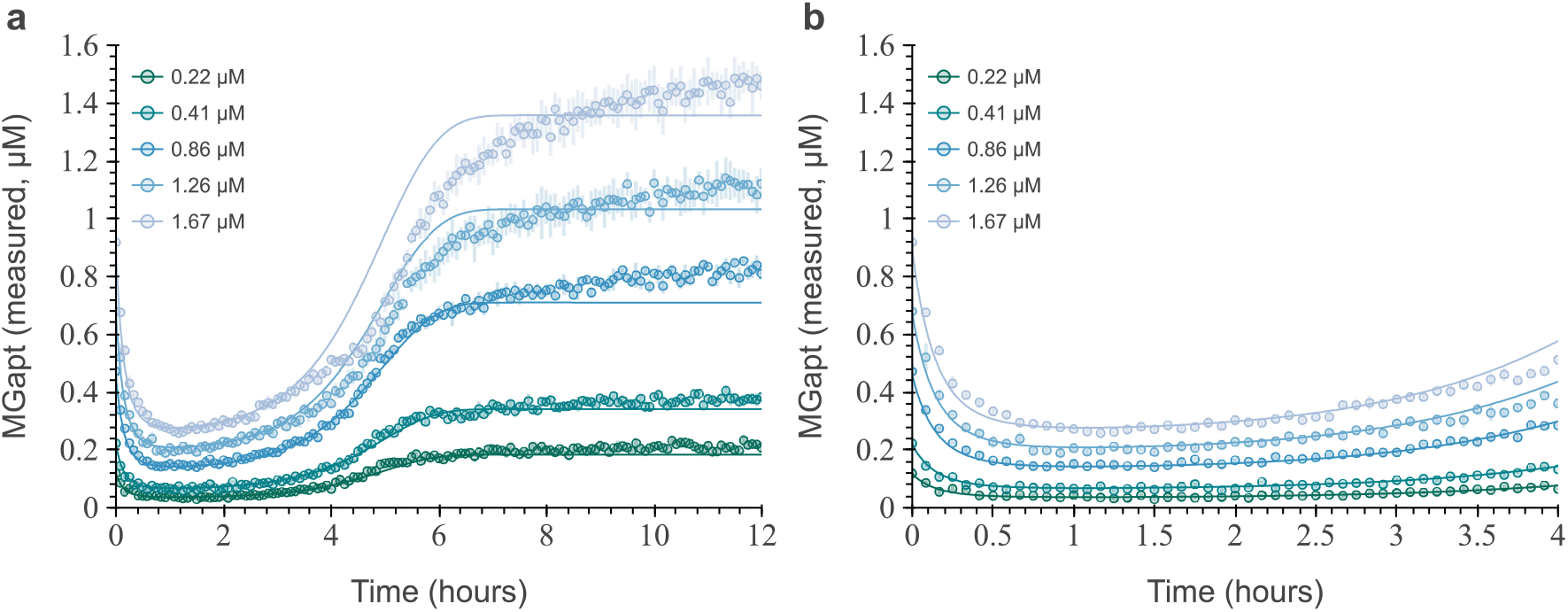
Modeled MGapt_measured_ at different initial RNA concentrations. The modeled MGapt_measured_ of BioTek at different RNA concentrations (solid line) overlaying with experimental data, three replicates (circles and error bars, of respective colors). (**a**) Full measured time of 12 hours. (**b**) Zoomed, reaction relevant time of 4 hours.

As seen in Figure 6a, the predicted MGapt_measured_ concentration remains reasonably accurate throughout the measured time for concentrations below 1 µm. The increased error above below 1 µm of RNA is most likely due to changes in DTT degradation or increased stability of the MGapt_altered_ state. Regardless, the model does recapitulate the increase of the measured MGapt concentration and its overall dynamics. Furthermore, as indicated in the manuals of commercial PURE systems, the reaction is generally monitored for a 2 h-4 h rather than 12 hour. Upon closer examination of the pertinent 4 h reaction window, highlighted in Figure 6b, it becomes evident that the model accurately predicts the measured MGapt_measured_. Moreover, the model consistently predicts the measured MGapt within an average of 10% margin of error (refer to Figure S5) within the initial 4 hour period for RNA concentration is greater than 1 µm, longer the recommended reading time. The accuracy and the capturing of the dynamics of measured MGapt fluorescence demonstrate that DTT can serve as a representative model for MGapt fluorescence, commonly employed in measuring RNA production. Moreover, the simulation can be used in reverse to calculate the total RNA in a system by calculating the total RNA ratio between measured and total to calculate back the amount of RNA, as illustrated in Figure S6.

## Conclusion

Our experiments reveal that the chemical makeup of commercial PURE systems influences measured MGapt concentration. Specifically, we observe a correlation between the chemical properties of DTT, given that DTT is absent in OnePot PURE but present in other commercial PURE systems. Though DTT and TCEP are effective reducing agents commonly used in biochemical research; DTT is more sensitive to oxidation, less stable, and more violative. We hypothesize that the concentration of DTT suppresses the fluorescence of MGapt by reversibly converting it into alternative forms with reduced fluorescence, which are indistinguishable and challenging for readers to account for. The degradation of DTT over time, driven by its chemical properties, enables MGapt to return to its fully fluorescent state, dynamics observed using purified RNA. We demonstrated that we could lengthen the suppression time by increasing the initial concentration of DTT in the expression of either transcription, translation expression, or both.

Moreover, given the inability to reduce the concentration of DTT in most commercial PURE systems, we introduce a model to predict the measured MGapt concentration by accounting for DTT’s impact on the state of MGapt. We identified a distribution of possible parameters in the model with the experimental data at one concentration of RNA for MGapt-UTR1-deGFP. We validated our model by accurately predicting the measured MGapt for the same RNA construct at five different concentrations within the relevant 2 hour period.

This model is essential in comprehending and accurately quantifying transcription within cell-free expression systems, specifically commercial PURE systems. Employing our methodology enables the construction of mathematical models based on the measured MGapt concentration and can be utilized to retroactively calculate the total quantity of RNA used in the PURE reaction. More importantly, the realization that MGapt can transition between various fluorescence states in cell-free expression systems suggests potential inaccuracies in our understanding and assessment of RNA production. While MGapt is widely and extensively utilized as an aptamer, this phenomenon may not be exclusive. Exploring other aptamers and monitoring fluorescence changes over time could prove beneficial in identifying additional chemical reactions or environmental factors that might influence the measured RNA concentration.

## Materials and Methods

### Computational Modeling and Simulations

In this paper, we build a deterministic model of two fluorescently different MGapt states driven by the concentration of DTT. This model is based on MGapt different states effects by organic solvents found Stead *et al*., Zhou *et al*., and Da Costa *et al*. [22, 15, 19]. We use the chemical reaction network (CRN) formalism to create the detailed mechanistic model using a CRN compiler called BioCRNpyler [30]. The computational model was developed to take a total RNA concentration of MGapt-UTR1-deGFP and predict the measured MGapt concentration using a BioTek plate reader. Our CRN model consists of five distinct species and seven total reactions.

The BioCRNpyler tool generates models in the Systems Biology Markup Language (SBML)[33], a standard format for biological modeling. These SBML files can be simulated using any compatible SBML simulator. In our case, we utilize the Bioscrape Python package [32] for simulation. Bioscrape converts the Chemical Reaction Network (CRN) model to ordinary differential equations (ODEs) and employs Python’s `odeint` to solve them based on specified initial conditions. Each reaction rate in the CRN is expressed using mass-action propensity [34] for this conversion. We opt for Bioscrape because it supports sensitivity analysis, Bayesian inference tools, and model simulations. Local sensitivity analysis is conducted for each SBML model to determine the sensitivity of measured species to all parameters over time. Subsequently, we identify the most sensitive parameters using experimental data. Parameter identification is achieved through a Bayesian inference algorithm implemented in Bioscrape, utilizing the emcee Python package [35]. By incorporating experimental data, we obtain probability distributions for each identified parameter through Bayesian inference. Model simulations, using parameter values sampled from these posterior probability distributions, are then compared against experimental data to assess model prediction quality. These posterior probability distributions also quantify the uncertainty in the data, highlighting a significant advantage of Bayesian inference methods.

Summarizing the computational analysis for the PURE transcription model, we identified five out of seven parameters using the experimental data. The final trained model predicts the BioTeK measurement of MGapt concentration over time for the RNA sequence of MGapt-UTR1-deGFP. All calculations and plotting were performed using the standard stack of Python packages – NumPy [36], SciPy [37], Pandas [38], Matplotlib [39], Bokeh [40], and Seaborn [41]. The simulation time for one RNA concentration is less than a tenth of a second on a personal computer running AMD Ryzen 7 4700U 2.0 GHz with 16GB of RAM.

### DNA constructs

The original DNA plasmids were procured from the Arbor Biosciences myTXTL Toolbox 2.0 plasmid collection [42], pTXTL-T7p14-mGapt and pTXTL-T7p14-deGFP. The deGFP protein is on a T7 promoter with a UTR1 ribosome binding site (RBS). Primers from Integrated DNA Technologies were employed to clone the MGapt between the promoter and the RBS site. Detailed information regarding the forward and reverse primers utilized can be found in the Supplementary Information Table S5.

Subsequently, both plasmids were initially transformed into JM109 cells and cultivated overnight on plates containing carbenicillin resistance at 37 °C. Colonies from each plate were selected and cultured in 4.5 mL LB medium with 4.5 µL of carbenicillin (100 mg*/*µL) overnight. Glycerol stocks of each plasmid were prepared by mixing 500 µL of liquid culture with an equal volume of 50% glycerol, while the remaining culture was miniprep using the Qiagen Miniprep Kit. Before running DNA in the cell-free reaction, all plasmids underwent an additional PCR purification step utilizing a QiaQuick column (Qiagen) to eliminate excess salt. The purified plasmids were then eluted and stored in nuclease-free water at 4 °C for short-term storage and at −20 °C for long-term storage.

### RNA constructs

Purified RNA was made using the original plasmid of pTXTL-T7p14-mGapt (pT7-MGapt-tT7) and the modified pTXTL-T7p14-deGFP (pT7-MGapt-UTR1-deGFP-tT7) plasmid. Purified plasmids were first amplified and linearized using Q5 Hot Start High-Fidelity DNA Polymerase (NEB); forward and reverse primers are listed in Table S5. Eight reactions of 125 µL, for a total volume of 1 mL were run in a thermocycler. The thermocycler was run for 30 cycles with an elongation time of 15 sec or 45 sec, respectively, and annealing temperature of 65 °C. The completed PCR reactions were then combined into three microcentrifuge tubes of 333 µL to which 33 µL sodium acetate (3 m) and 1 mL Ethanol (100%) was added. All three tubes were placed at −80 °C for 20 min and then centrifuged at 16 000 xg for 30 min at 4 °C to remove all traces of ethanol. The precipitated DNA was then resuspended in 50 µL of nuclease-free water, combined, and stored at 4 °C.

Using the HiScribe T7 High Yield RNA Synthesis Kit (NEB) 1 µg of the amplified and linearized DNA construct was transcribed in an 80 µL reaction volume and incubated overnight at 37 °C. Before the RNA purification procedure, the transcription reaction underwent DNAse I treatment for 15 min at 37 °C to remove the linear DNA template. Ethanol (100%) was then added to achieve a final concentration of 35%, to precipitating the RNA. Next, using PureLink RNA Mini Kit (Invitrogen), RNA was isolated following the protocol for ‘Purification of RNA from liquid samples’ starting from Step 2-Bind RNA. The yield and quality of the RNA samples were analyzed using the NanoDrop2000c measuring at UV absorbance at 260 nm. The purified RNA was flash frozen in liquid nitrogen and stored at −80 °C.

### PURE reactions and fluorescence measurements

The PURE reactions were prepared according to the NEB PURExpress (E6800) protocol, adapted for a 10 µL reaction volume, and incubated in a 384-well plate (Nunc) at 37 °C. A concentration of 5 nm DNA was utilized, unless otherwise specified, along with 8 units of RNAse inhibitor (NEB), and 10 µm malachite green oxalate added to each reaction. The RNA concentrations varied and were specified in the corresponding experimental setup description. DNA and RNA were added using an Echo 525 Acoustic Liquid Handler and final concentrations were recalculated based on dispensed volumes. The remaining components were made into a master mix with 5% excess, added to a 384-well plate using a repeater pipette.

Fluorescence readings were obtained using a BioTek Synergy H1 plate reader (BioTek) at 3-minute intervals for 12 hours at 37 °C. Excitation/emission wavelengths were set at 610*/*650 nm for MGapt with a gain of 150. All samples were analyzed using the same plate reader. For RNA calibrations, relative fluorescence units (RFUs) were converted to µm of RNA using purchased MGapt from IDT. Dilutions of MGapt (rArCrUrGrGrArUrCrCrCrGrArCrUrGrGrCrGrArGrArGrCrCrArGrGrUrArArCrGrArArUrGrGrArUrCrCrArArU) in PBS were run, and multiple reads per concentration were averaged to make a calibration plot, Figure S2.

## Acknowledgments

We thank Zachary Martinez for their encouragement during the start of this project. We sincerely appreciate Manisha Kapasiawala, Dr. Yan Zhang, and Dr. John Marken for their invaluable insights and thought-provoking discussions on the experiments and modeling framework throughout the project. We would also like to acknowledge the contributions of Dr. Ayush Pandey, who provided assistance and feedback on the proposed model’s modeling, analysis, and inference. Their involvement has been indispensable to its completion.

Research supported by the National Science Foundation award number 2152267. The computations presented here were conducted in the Resnick High Performance Computing Center, a facility supported by the Resnick Sustainability Institute at the California Institute of Technology, Pasadena, CA, USA.

