## Supplementary Information for "Impact of Chemical Dynamics of Commercial PURE Systems on Malachite Green Aptamer Fluorescence"

Zoila Jurado[1,*] and Richard M. Murray[1]

[1]Division of Engineering and Applied Science, California Institute of Technology, Pasadena, CA

March 26, 2024

### 1 Comparison between PURE and OnePot

The probable chemical composition of PURExpress, based on concentrations published by Shimizu *et al.* [1], and OnePot PURE, based on concentrations published by Grasemann *et al.* [2].

**Table S1:** Reported chemical composition of PURE systems

| Chemical | PURExpress | OnePot PURE |
|---|---|---|
| Magnesium acetate | 9 mM | 11.8 mM |
| Potassium phosphate | 5 mM | - |
| Potassium glutamate | 95 mM | 100 mM |
| Ammonium chloride | 5 mM | - |
| Calcium chloride | 0.5 mM | - |
| Spermidine | 1 mM | 2 mM |
| Creatine phosphate | 10 mM | 20 mM |
| Putrescine | 8 mM | - |
| Dithiothreitol (DTT) | 1 mM | - |
| Tris(2-carboxyethyl)phosphine (TCEP) | - | 1 mM |

#### 2 pH of the PURE system using SNARF-5F

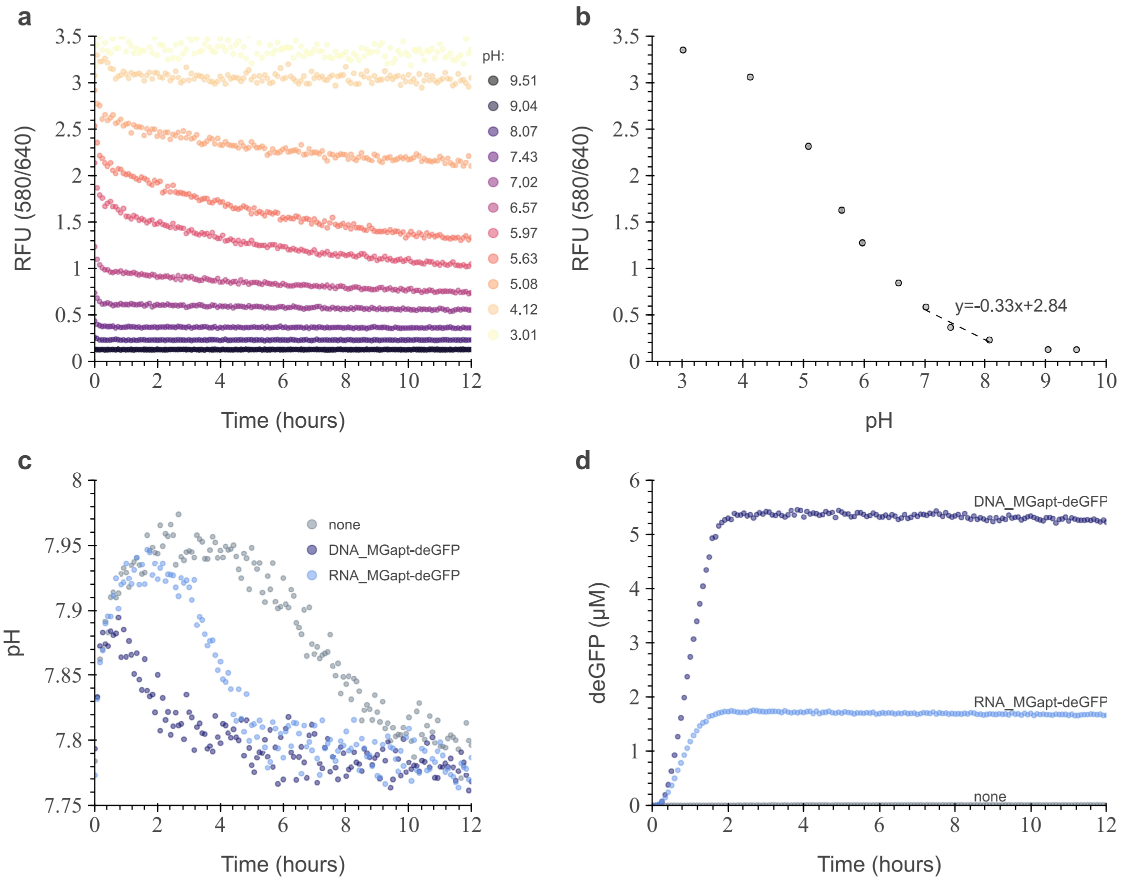


**Figure S1: Measurement of pH of the PURExpress over time. (a)** The pH was measured using SNARF-5F. The ratio of 580 RFU over 640 RFU data of PBS solutions, at respective pH, was used for pH calculations. **(b)** Calibration curve for pH calculations. Each point represents the average RFU(580/640) over the 12-hour read. A linear regression line was fitted between pH 7.02 and pH 8.07. **(c)** The pH of PURExpress under three different expression conditions: none-without DNA (gray circles), DNA plasmid of pT7-MGapt-UTR1-deGFP-tT7 (light blue circles), and RNA of pT7-MGapt-UTR1-deGFP (dark blue circles). **(d)** Production of deGFP over pH assay.

### 3 MGaptamer RNA Calibration

#### Traditional RNA Calibrations

The fluorescence calibration curve for MGaptamer was generated using single-stranded RNA purchased from Integrated DNA Technologies (IDT), rArCrUrGrGrArUrCrCrCrGrArCrUrGrGrCrGrArGrArGrCrCrArGrGrUrArArCrGrArArUrGrGrArUrCrCrArArU. The sample arrived lyophilized in tube and weighed 64.5 nmol (0.92 mg). To achieve the concentration of 100 µM 645 µL of nuclease-free water (NFW) was added. The sample was vortexed for several minutes and heated to 55°C for 5 min before vortexing again. The stock concentration was measured by a Nanodrop 2000c before serial dilutions in 1X PBS. Next, respective dilutions were deposited onto the bottom of a Nunc 384 well plate using an Echo 525 Acoustic Liquid Handler. Each well contained a total volume of 10 µL with four technical replicates containing 10 µM of Malachite Green dye. The Nunc 384 well plate was read using a BioTeK H1MF plate reader at 37 °C and at 610/650 (Ex/Em) and gain 150. Each point on each calibration curve represents the average of 20 points, and four replicates were read over 10 minutes at 2.5-minute intervals to generate 5 points per replicate. The points were all background-subtracted from the negative control such that the 0 µM samples had zero fluorescence. Points were fit using linear regression and were not forced to go through the origin. Fits for each calibration curve are indicated in Fig. S2.

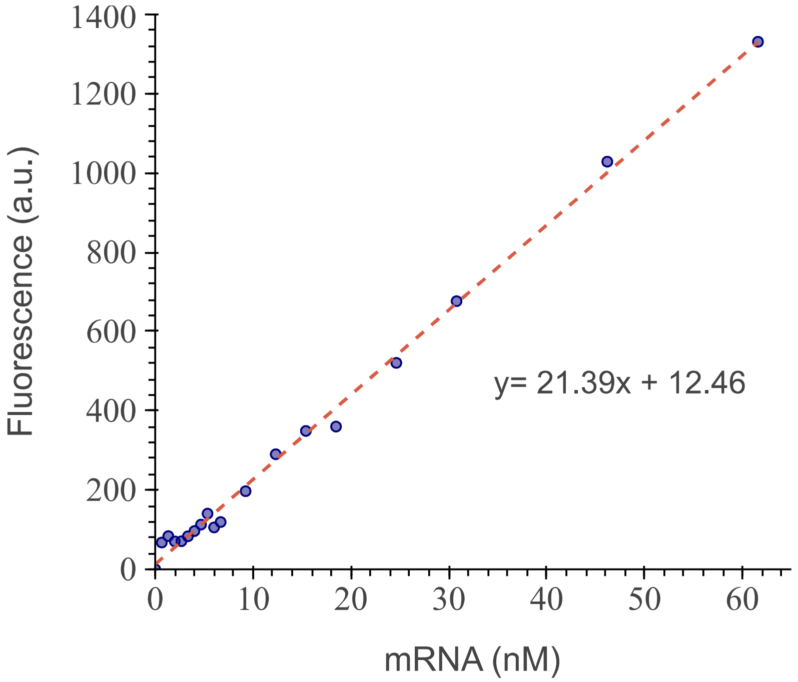


**Figure S2: Traditional MGapt calibration curve.** The fluorescence calibration curve for MGapt used to convert RFU to µM.

### 4 Effects of Buffer on Measured MGapt Concentration

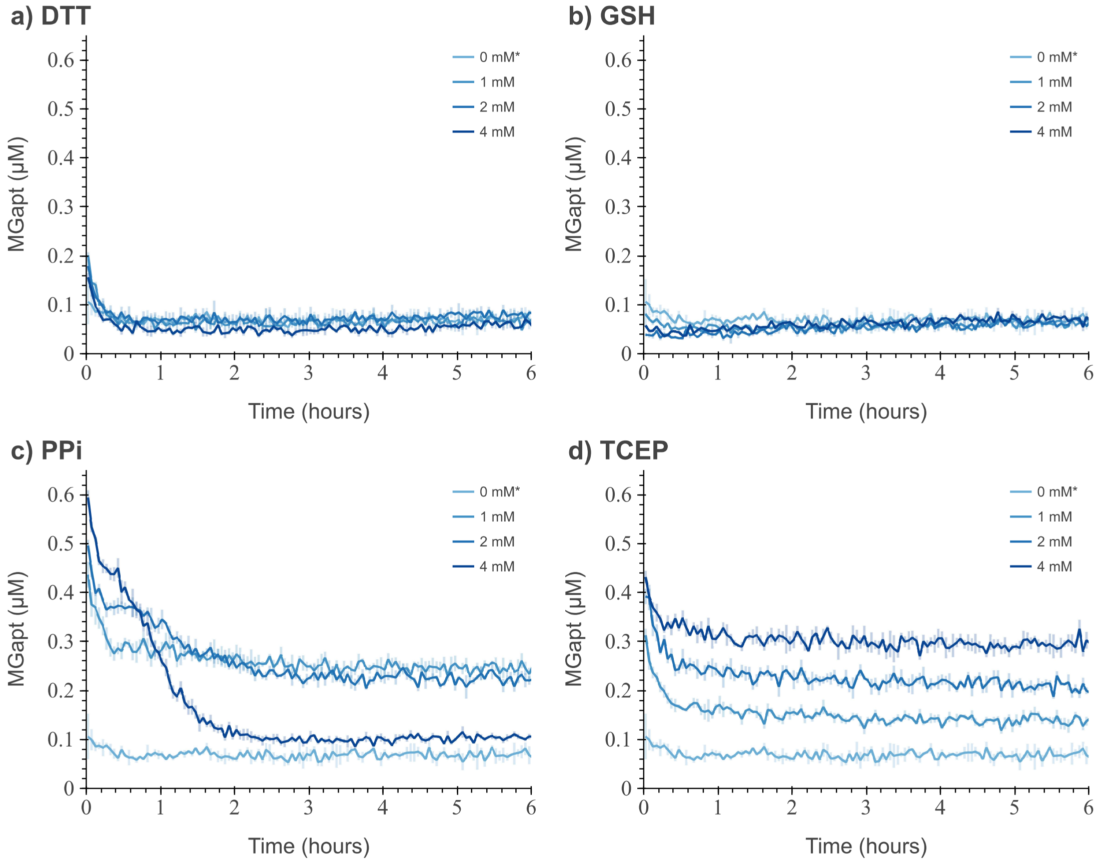


**Figure S3: MGapt measurement in different buffers.** Measurements of MGapt concentration of purified RNA for MGapt-UTR1-deGFP at 0.51 µM in different reducing agents and waste chemicals over four hours. Each subplot is titled with the respective buffer used: (**a**) dithiothreitol (DTT), (**b**) glutathione (GSH), (**c**) pyrophosphate (PPi) and (**d**) tris(2-carboxyethyl)phosphine (TCEP). The chemical concentrations are in different shades of blue, respectively. The plot shows the average of the three replicated with error bars; negative control without MGapt was subtracted.

### 5 Measured MGapt concentration dynamics in PURExpress

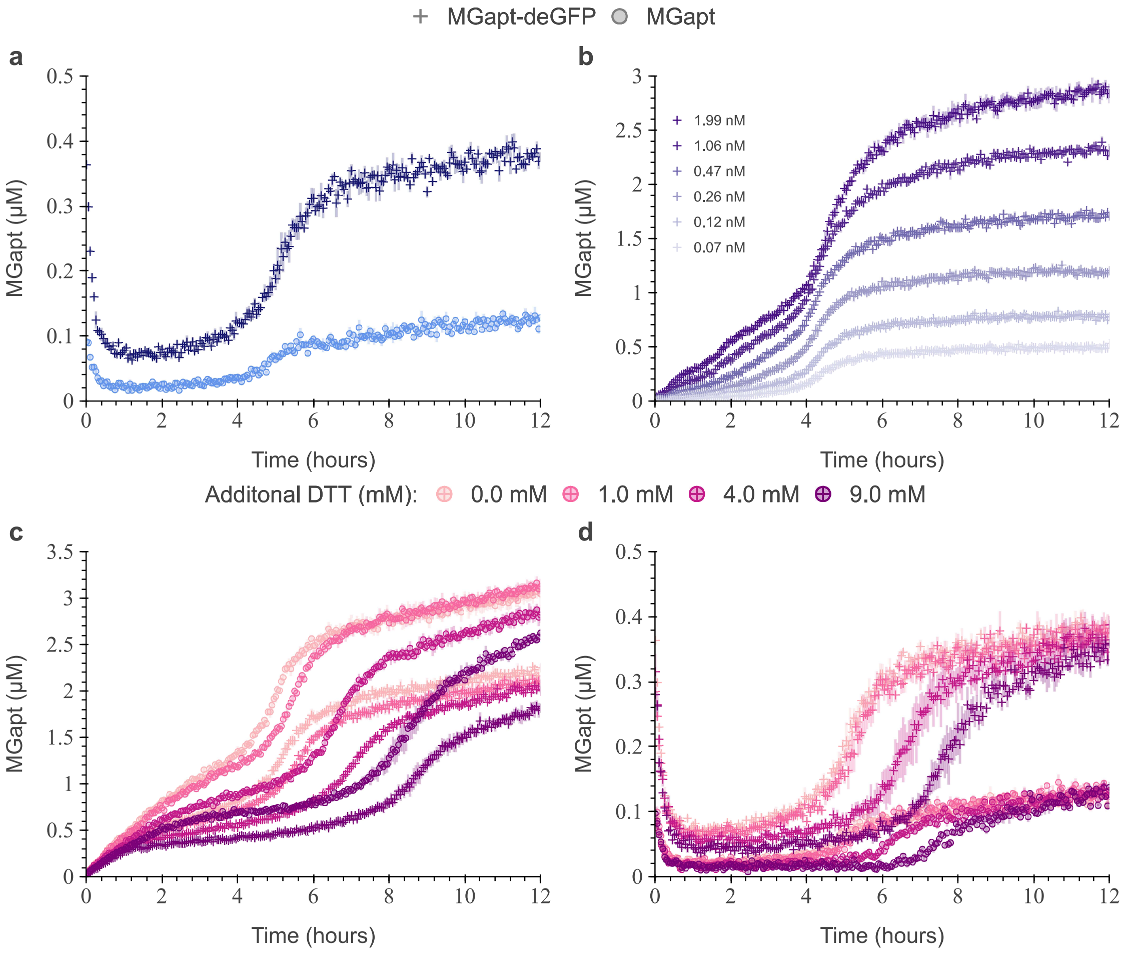


**Figure S4: Measured MGapt concentration dynamics in PURExpress.** PURExpress reactions of 10 µL with three technical replicates containing 10 µM of Malachite Green dye and 8 units of RNAse inhibitor. (**a**) Starting from purified RNA of MGapt at 0.53 µM (light blue crosses) and MGapt-UTR1-deGFP at 0.49 µM (dark blue circles). (**b**) Expressing plasmid pT7-MGapt-UTR1-deGFP-tT7 at various DNA concentrations. (**c**) Expressing of plasmid pT7-MGapt-UTR1-deGFP-tT7 at 4.93 nM and pT7-MGapt-tT7 at 5.03 nM with increasing DTT concentration. (**d**) Starting from purified RNA of MGapt-UTR1-deGFP at 0.49 µM and MGapt at 0.53 µM with increasing DTT concentration.

### 6 Initial Conditions

The model's initial conditions parameters depend on the total amount of RNA added by the user, where $\sum$MGapt is equal to the total RNA containing MGapt added. Initial conditions account for the formation of $\text{MGapt}_{\text{bound}}$ and the $\text{MGapt}_{\text{altered}}$ before starting the BioTek measuring. In the model, we make the calculation that 45 % and 55 % of the $\sum$MGapt are in the two different states of $\text{MGapt}_{\text{bound}}$ and $\text{MGapt}_{\text{altered}}$, respectively. Therefore, $\text{MGapt}_{\text{t=0}}$ is zero and $\text{MGapt}_{\text{bound,t=0}} = 0.45 \sum \text{MGapt}$, $\text{MGapt}_{\text{altered,t=0}} = 0.55 \sum \text{MGapt}$, and $\text{MG}_{\text{dye,t=0}} = \text{MG}_{\text{dye,total}} - \sum \text{MGapt}$.

#### 6.1 Initial Conditions for Analysis and Training Model

In this simulation of the model, the $\sum$MGapt is 0.22 µM. The initial conditions values for the MGapt-DTT model used in analysis and training are given in Table S2.

**Table S2:** MGapt-DTT model initial conditions.

| Species | Value | Unit |
|---|---|---|
| MGapt | 0 | µM |
| $\text{MGapt}_{\text{bound}}$ | 0.099 | µM |
| $\text{MGapt}_{\text{altered}}$ | 0.121 | µM |
| DTT | 1000 | µM |
| $\text{MG}_{\text{dye}}$ | 9.78 | µM |

#### 6.2 Initial Conditions for Validation Test

The model's initial conditions parameters depend on the total amount of RNA added by the user. In the simulation, to validate the model, $\sum$MGapt is varied. The initial conditions values for $\text{MGapt}_{\text{bound,t=0}}$, $\text{MGapt}_{\text{altered,t=0}}$, and $\text{MG}_{\text{dye,t=0}}$ used in validating the model are provided in Table S3. The initial conditions for MGapt and DTT are not affected by the change of RNA added by the user.

**Table S3:** MGapt-DTT model initial conditions.

| $\sum$MGapt, total RNA added (µM) | $\text{MGapt}_{\text{bound}}$ (µM) | $\text{MGapt}_{\text{altered}}$ (µM) | $\text{MG}_{\text{dye}}$ (µM) |
|---|---|---|---|
| 0.41 | 0.1845 | 0.2255 | 9.59 |
| 0.86 | 0.387 | 0.473 | 9.14 |
| 1.26 | 0.567 | 0.693 | 8.74 |
| 1.67 | 0.7515 | 0.9185 | 8.33 |

### 7 Parameters Values

The parameter values for the MGapt-DTT model are given in Table S4. The initial chemical reaction rates of the MGapt-DTT model were first-hand-tuned and then run through Bayesian inference tools in Bioscrape [3].

**Table S4:** MGapt-DTT model parameters values.

| Parameter | Description | Value | Unit |
|---|---|---|---|
| $k_{\mathrm{MGapt}}$ | Formation of $\mathrm{MGapt_{bound}}$ though the binding of $\mathrm{MG_{dye}}$ and MGapt | $9.5 \times 10^{-4}$ | $\mu\mathrm{M}^{-1}\,\mathrm{h}^{-1}$ |
| $k_{\mathrm{b_{DTT}}}$ | Binding of DTT to $\mathrm{MGapt_{bound}}$ to form $\mathrm{MGapt_{altered}}$ | $2.0 \times 10^{-6}$ | $\mu\mathrm{M}^{-1}\,\mathrm{h}^{-1}$ |
| $k_{\mathrm{u_{DTT}}}$ | Unbinding of DTT from $\mathrm{MGapt_{altered}}$ | $8.0 \times 10^{-6}$ | $\mathrm{h}^{-1}$ |
| $k_{\mathrm{u_{MGapt}}}$ | Unbinding of DTT and $\mathrm{MG_{dye}}$ from $\mathrm{MGapt_{altered}}$ | 0.0675 | $\mathrm{h}^{-1}$ |
| $k_{\mathrm{b_{MGapt}}}$ | Rebinding of DTT and $\mathrm{MG_{dye}}$ and MGapt to reform $\mathrm{MGapt_{altered}}$ | 0.0762 | $\mu\mathrm{M}^{-2}\,\mathrm{h}^{-1}$ |
| $k_{\mathrm{RNA_{deg}}}$ | Degradation of MGapt | $3.2 \times 10^{-4}$ | $\mathrm{h}^{-1}$ |
| $k_{\mathrm{DTT_{deg}}}$ | Degradation of DTT | $3.2 \times 10^{-4}$ | $\mathrm{h}^{-1}$ |

The model's parameter value depends only on the amount of RNA added and the DTT concentration. Other auxiliary reactions, which may play a role in DTT interaction with MGapt, were not incorporated.

### 8 Error of BioCRNpyler Model

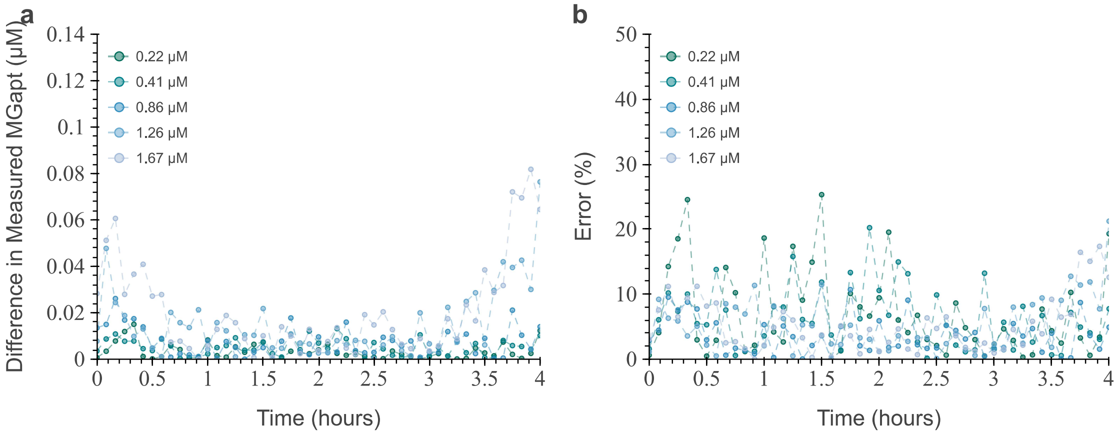


**Figure S5: Error of proposed BioCRNpyler model.** (**a**) Absolute error between the BioCRNpyler model and experimental data for different initial RNA concentrations. (**b**) Percent error between the BioCRNpyler model and experimental data for different initial RNA concentrations.The model compared to the mean of the experimental results for the respective RNA concentrations.

### 9 Dynamic MGapt Calibration

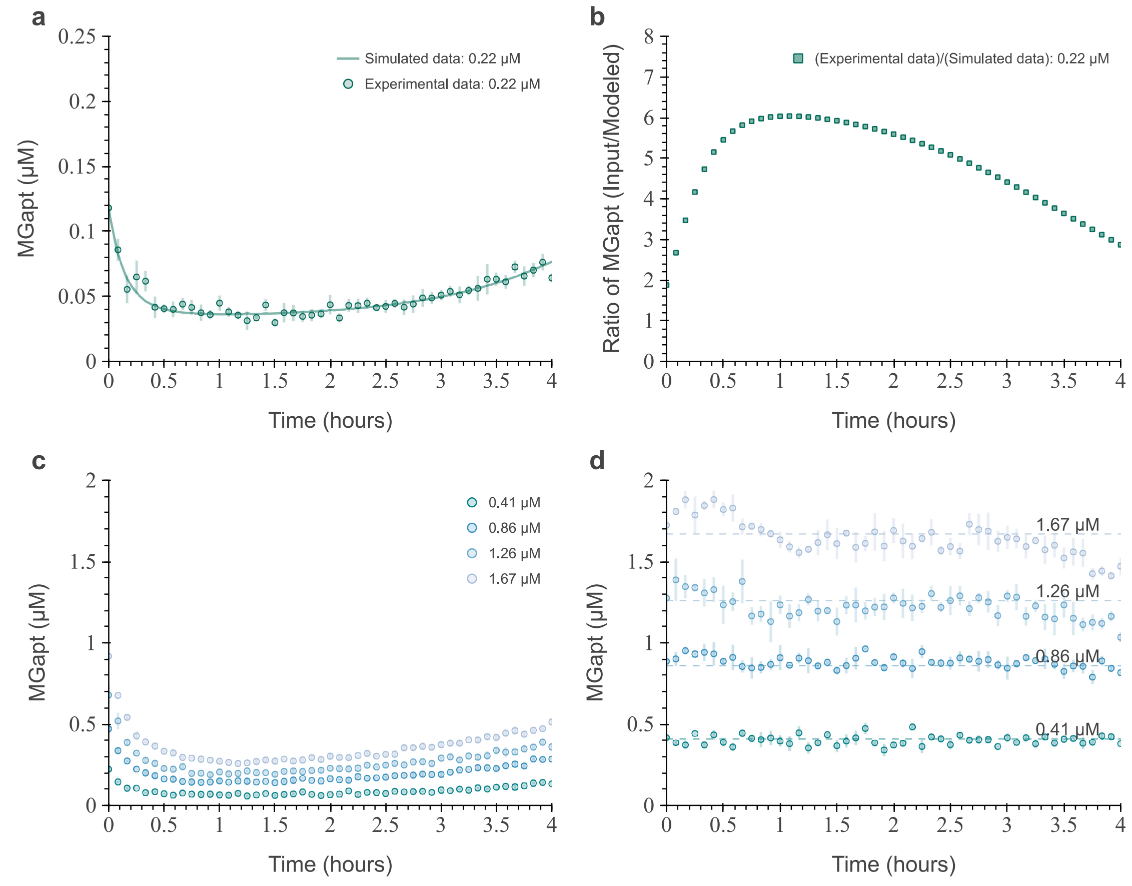


**Figure S6: Dynamic calibrations of MGapt.** (**a**) Measured MGapt concentration of 0.22 μM of RNA for MGapt-UTR1-deGFP (circles with error bars) and simulated MGapt measurements (solid line). (**b**) Proposed dynamic calibration curve for MGapt concentration measurements from data shown in (a). (**c**) Measured MGapt of RNA, MGapt-UTR1-deGFP, at concentrations: 0.41 μM, 0.86 μM, 1.26 μM,and 1.67 μM (circles with error bars, at respective color). (**d**) Calibrated experimental data from (c) with dynamic calibration curve in (b) (circles with error bars, at respective color), overlayed with the RNA concentration used (dashed lines).

### 10 Primers

**Table S5:** Primers used to clone MGapt, GGGATCCCGACTGGCGAGAGCCAGGTAACGAATGGATC, into pTXTL-T7p14-deGFP DNA originally obtained from myTXTL [4] and to linearize DNA for RNA purifications. The bold text identifies the binding region of the plasmid.

| Name | Seq | Purpose |
|---|---|---|
| pT7_MGapt_FOR | GAGCCAGGTAACGAATG GATCCAAT**AATTTTGT TTAACTTTAAGAAGG AGATATACCATG** | Cloning in MGapt to pTXTL-T7p14-deGFP |
| pT7_MGapt_REV | ATTGGATCCATTCGTTA CCTGGCTCTCGCCAGTC GGGATCCC**TCTAGAGG GAAACCGTTG** | Cloning in MGapt to pTXTL-T7p14-deGFP |
| pPCR_MGapt_FOR | **GTGATGTCGGCGATA TAGGC** | Linearize pTXTL-T7p14-mGapt |
| pPCR_MGapt_REV | **CACTATCGACTACGC GATCATG** | Linearize pTXTL-T7p14-mGapt |
| pPCR_MGapt-UTR1-deGFP_FOR | **GCGTAGAGGATCGAG ATCTCGATC** | Linearize modified pTXTL-T7p14-deGFP |
| pPCR_MGapt-UTR1-deGFP_REV | **CTATCGACTACGCGA TCATGGC** | Linearize modified pTXTL-T7p14-deGFP |

[2] Laura Grasemann, Barbora Lavickova, M. Carolina Elizondo-Cantú, and Sebastian J. Maerkl. OnePot PURE cell-free system. *JoVE*, (172):e62625, 2021.

[3] Ayush Pandey, William Poole, Anandh Swaminathan, Victoria Hsiao, and Richard M. Murray. Fast and flexible simulation and parameter estimation for synthetic biology using bioscrape. *Journal of Open Source Software*, 8(83):5057, 2023.

[4] Arbor Biosciences. *myTXTL T7 Expression Kit.* Michigan, United States of America, 2019.
