## Supplementary Data Files for "Impact of Chemical Dynamics of Commercial PURE Systems on Malachite Green Aptamer Fluorescence": DTTvMGapt_Figures_Final.html

March 26, 2024

[1] Division of Engineering and Applied Science, California Institute of Technology, Pasadena, CA \

This notebook was used to plot all figures depicted in the *Impact of Chemical Dynamics of Commercial PURE Systems on Malachite Green Aptamer Fluorescence* and supplementary information.

### Importing required packages and definitions¶

In [1]:

```
#workhorses
import numpy as np
import pandas as pd
from scipy.signal import savgol_filter
import math
from scipy import stats
import seaborn as sns; sns.set()
import random 
import holoviews as hv
import matplotlib.pyplot as plt
from matplotlib.colors import LogNorm, Normalize
from matplotlib.ticker import MaxNLocator
from matplotlib.colors import LinearSegmentedColormap
import seaborn as sns; sns.set_theme()
%matplotlib inline

# Bokeh
import bokeh
import bokeh.io
import bokeh.plotting
bokeh.io.output_notebook()
hv.extension('bokeh')
# Modules needed from Bokeh.
from bokeh.models import ColumnDataSource, FactorRange
from bokeh.models import LinearAxis, Range1d
from sklearn.metrics import r2_score
from bokeh.io import export_svgs, export_svg
from bokeh.models import Title
from bokeh.plotting import gridplot,figure, output_file, show
from bokeh.models.glyphs import Text
from bokeh.transform import factor_cmap
from bokeh.themes import Theme
from bokeh.models import Legend, LegendItem
from bokeh.models import Label

#From bioscrape
from bioscrape.simulator import py_simulate_model, ModelCSimInterface, DeterministicSimulator
from bioscrape.types import Model

#Get directory
import os
directory0 = os.getcwd()
directory=directory0+'/MGapt_deGFP_Exp_PURE_Data'
```

Loading BokehJS ...

#### Definitions¶

In [2]:

```
def create_custom_plot(title_text, x_max=8,y_max=2, xname='Time (hours)', yname='MGapt (μM)',height=400, width=500):
    custom_plot = figure(
        toolbar_location='right',
        outline_line_color=None,
        min_border_right=10,
        height=height,
        width=width,)

    custom_plot.title.text = title_text
    custom_plot.xaxis.axis_label = xname
    custom_plot.yaxis.axis_label = yname
    custom_plot.y_range = Range1d(0, y_max)
    custom_plot.x_range = Range1d(0, x_max)
    custom_plot.outline_line_color = None

    # custom_plot.yaxis
    custom_plot.ygrid.visible = False
    custom_plot.yaxis.axis_label_text_font_size = '15pt'
    custom_plot.yaxis.major_label_text_font_size = '15pt'
    custom_plot.yaxis.major_label_text_font = 'Work Sans'
    custom_plot.yaxis.axis_label_standoff = 15
    custom_plot.yaxis.axis_label_text_font_style = 'normal'

    # custom_plot.xaxis
    custom_plot.xgrid.visible = False
    custom_plot.xaxis.axis_label_text_font_size = '15pt'
    custom_plot.xaxis.major_label_text_font_size = '15pt'
    custom_plot.xaxis.major_label_text_font = 'Work Sans'
    custom_plot.xaxis.axis_label_standoff = 15
    custom_plot.xaxis.axis_label_text_font_style = 'normal'

    # custom_plot.title
    custom_plot.title.text_font_size = '18pt'
    custom_plot.title.align = "left"
    custom_plot.title.offset=-70

    return custom_plot
```

In [3]:

```
def Circle_wErrorPlot(plot, DF, ind, color='black',cal=False, marker="circle",size=5,):
    
    if cal==True:
        #Data from experiments
        plot.scatter(
            x=DF['Time']/3600, y=calibrateBiotek4(DF['value_ave']),  marker=marker,color= color, size=size, fill_alpha=0.2,
            legend_label= ind)
        # Add error bars
        plot.segment(
            x0=DF['Time']/3600, y0=calibrateBiotek4(DF['error_low']),
            x1=DF['Time']/3600, y1=calibrateBiotek4(DF['error_high']),
            line_width=2, color= color, line_alpha=.25)
    else:
        plot.scatter(
            x=DF['Time']/3600, y=(DF['value_ave']),  marker=marker,color= color, size=size, fill_alpha=0.2,
            legend_label= ind,)
        # Add error bars
        plot.segment(
            x0=DF['Time']/3600, y0=(DF['error_low']),
            x1=DF['Time']/3600, y1=(DF['error_high']),
            line_width=2, color= color, line_alpha=.25)
    
    plot.legend.location="top_left"
    plot.legend.click_policy="hide"
    plot.legend.border_line_color = None
    plot.legend.background_fill_color = None
    return(plot)
```

In [4]:

```
def Cal_avesNsems(df, DF_neg=None, norm=False, yname='value',xname='Time'):
    num=len(df['well'].unique())
    length=int(len(df)/num)
    coln=array_repeats[0:num]
    arr = df[yname].values.copy()
    arr.resize(num,length)
    DF=pd.DataFrame(arr).T
    
    #Subtract the negative control
    if DF_neg is None:
        pass
    else:
        for n in coln:
            DF[n]=DF[n]-DF_neg['value_ave']
    
    #Normalize data if needed    
    if norm==False:
        pass
    else:
        for n in coln:
            DF[n]=DF[n]/DF[n].max()
            
    DF['value_ave']=DF.iloc[:, coln].mean(axis=1)    
    DF['sem']=stats.sem(DF.iloc[:,coln].T)
    DF[xname]=df[xname].reset_index(drop=True)[0:length]*60
    # Add error bars to the DataFrame
    DF['error_low'] = DF['value_ave'] - DF['sem']
    DF['error_high'] = DF['value_ave'] + DF['sem']
    return(DF)
```

In [5]:

```
def Cal_avesNsems2(df, DF_neg=None):
    
    #Calculating the measured MGapt
    total_MGapt=pd.DataFrame(columns=['mRNA0', 'mRNA1','mRNA2'])
    
    for name in total_MGapt.columns:
        total_MGapt[name]=df[name]-DF_neg['mRNA0']

    total_MGapt['time']=df['time']
    
    
    
    DF=total_MGapt.copy()
    DF['value_ave']=DF.iloc[:, [0,1,2]].mean(axis=1)
    DF['sem']=stats.sem(DF.iloc[:, [0,1,2]].T)
    DF['Time']=total_MGapt['time']

    # Add error bars to the DataFrame
    DF['error_low'] = DF['value_ave'] - DF['sem']
    DF['error_high'] = DF['value_ave'] + DF['sem']
    
    return(DF)
```

In [6]:

```
def calibrateBiotek4(df):
    cal_data=(df-12.46)/21.39/1000
    return(cal_data)
```

In [7]:

```
def updateLegend(plot, title="", location='top_left',):
    plot.legend.location=location
    plot.legend.click_policy="hide"
    # Remove the box around the legend
    plot.legend.border_line_color = None
    # Add a legend and set its title
    plot.legend.title = title
    plot.legend.title_text_font_style = "normal"
    # plot.legend.title_text_font_size = '15pt'
    # plot.add_layout(plot.legend[0],layout_loc)
    return(plot)
```

#### General colors and dictionary¶

In [8]:

```
## Define array for replicates
array_repeats = [0,1,2,3,4,5,6,7,8,9,10]
## Colors used:
colorBlue= ['#6baed6' , '#4292c6',"#2171b5",'#084594']
colorPurples=['#dadaeb','#bcbddc','#9e9ac8','#756bb1','#54278f','#4a1486']
colorN= {'mRNA0':"lightblue", 'mRNA1':"cornflowerblue",'mRNA2':"steelblue"}
colorReds=['#fcae91','#fb6a4a','#de2d26','#a50f15']
colorDTT=['#7a0177','#c51b8a','#f768a1','#fbb4b9',]
colorsmRNA=['#016c59', '#02818a','#3690c0','#67a9cf','#a6bddb','#d0d1e6',]
colorspH= bokeh.palettes.Magma[11]
## Dictionary for plotting
markers={'MGapt':'circle','MGapt-deGFP':'cross'}
size={'circle':5,'cross':8}
colorNA= {'none':"slategray",'MGapt-deGFP':"midnightblue", 'MGapt':"cornflowerblue",}
colorSNARF= {'none':"slategray",'DNA_MGapt-deGFP':"midnightblue", 'RNA_MGapt-deGFP':"cornflowerblue",}
```

### **Figure 1: Comparing PURExpress and OnePot results**¶

#### a) PURExpress¶

##### Get data¶

In [9]:

```
folder= '/2024.01.12_PURExpress_exp/'
filename = '2024.01.12_PURExpress_repeat_exp.xlsx'
data_dict = pd.read_excel(directory + folder+filename, sheet_name=None, engine='openpyxl')
sheets_tidy= [x for x in data_dict.keys() if '_tidy' in x]
df_mgapt=pd.read_csv(directory + folder+'/'+ sheets_tidy[1]+'.csv')
```

```
C:\Users\zoila\anaconda3\envs\python38\lib\site-packages\openpyxl\worksheet\_reader.py:329: UserWarning: Unknown extension is not supported and will be removed
  warn(msg)
C:\Users\zoila\anaconda3\envs\python38\lib\site-packages\openpyxl\worksheet\_reader.py:329: UserWarning: Unknown extension is not supported and will be removed
  warn(msg)
```

##### Index and plot¶

In [10]:

```
neg=df_mgapt[df_mgapt['construct']=='none']
mgapt=df_mgapt[df_mgapt['construct']=='MGapt']
mgapt_degfp=df_mgapt[df_mgapt['construct']=='MGapt-deGFP']

DF_Neg=Cal_avesNsems(neg)
DF_MGapt=Cal_avesNsems(mgapt,DF_neg=DF_Neg)
DF_MGaptdeGFP=Cal_avesNsems(mgapt_degfp,DF_neg=DF_Neg)
```

In [11]:

```
#A: MGapt measurements PURE
pNEB=create_custom_plot('a', x_max=12, y_max=3,)
Circle_wErrorPlot(pNEB, DF_MGapt, 'MGapt', "cornflowerblue",cal=True)
Circle_wErrorPlot(pNEB, DF_MGaptdeGFP,'MGapt-UTR1-deGFP',"midnightblue",cal=True)
pNEB.legend.location="bottom_right"
pNEB.legend.click_policy="hide"
# Remove the box around the legend
pNEB.legend.border_line_color = None
bokeh.io.show(pNEB)
```

#### b) OnePot PURE¶

##### Get data¶

In [12]:

```
folder= '/2024.01.08_OnePotPURE_exp/'
filename = '2024.01.08_OnePotPURE_MGaptMeasurements.xlsx'
data_dict = pd.read_excel(directory + folder+ filename, sheet_name=None, engine='openpyxl')
sheets_tidy= [x for x in data_dict.keys() if '_tidy' in x]
df_mgapt=pd.read_csv(directory + folder+'/'+ sheets_tidy[1]+'.csv')
```

##### Index and plot¶

In [13]:

```
neg=df_mgapt[df_mgapt['construct']=='None']
mgapt=df_mgapt[df_mgapt['construct']=='MGapt']
mgapt2=df_mgapt[df_mgapt['construct']=='MGapt-deGFP']

DF_Neg=Cal_avesNsems(neg)
DF_MGapt=Cal_avesNsems(mgapt,DF_neg=DF_Neg)
DF_MGaptdeGFP=Cal_avesNsems(mgapt2,DF_neg=DF_Neg)
```

In [14]:

```
#B: MGapt measurements PURE
pOnepot=create_custom_plot('b', x_max=8, y_max=1)
Circle_wErrorPlot(pOnepot, DF_MGapt, 'MGapt', "cornflowerblue",cal=True)
Circle_wErrorPlot(pOnepot, DF_MGaptdeGFP,'MGapt-UTR1-deGFP',"midnightblue",cal=True)
pOnepot.legend.location="bottom_right"
pOnepot.legend.click_policy="hide"
# Remove the box around the legend
pOnepot.legend.border_line_color = None
bokeh.io.show(pOnepot)
```

#### Plot as Grid and save as .svg¶

In [15]:

```
# using grid layout on plots
pCFPS=gridplot([[pNEB, pOnepot]], width=500, height=400)

count=1
for pic in [pNEB, pOnepot]:
    pic.output_backend = "svg"
    export_svgs(pic, filename = 'Figures_DTTvMGapt/Fig1_pCFPS' + str(count) + '.svg',width=500, height=400)
    count+=1
export_svgs(pCFPS, filename = 'Figures_DTTvMGapt/Fig1_pCFPS.svg')
show(pCFPS)
```

### **Figure 2: Comparing MGapt measurement in different buffers**¶

#### **Scatter plots**¶

##### Definitions¶

In [16]:

```
BufferTitles= {'DTT': 'a) DTT','GSH':'b) GSH','Ppi': 'c) PPi','TCEP': 'd) TCEP'}
```

In [17]:

```
def extract_numerical_part(item):
    return float(re.search(r'\d+(\.\d+)?', item).group())
```

In [18]:

```
def BufferPlots(buffer, data,idxNC):
    buffer_mean=[]
    buffer_sem=[]
    
    idxBuffer=data['buffer']==buffer
    df_Buffer=data[idxBuffer | idxNC]
    df_Buffer_naNeg=df_Buffer[df_Buffer['mrna']== 'no']
    df_Buffer_naPos=df_Buffer[df_Buffer['mrna']!= 'no']
    
    pBuffer=create_custom_plot(BufferTitles[buffer], x_max=6, y_max=.65)
    i=0
    # original_array=df_Buffer_naNeg['buffer_conc'].unique()
    sorted_array = ['0 mM','1 mM','2 mM','4 mM',] #sorted(original_array, key=extract_numerical_part)
    for buffer_conc in sorted_array:
        pos=df_Buffer_naPos[df_Buffer_naPos['buffer_conc']== buffer_conc]
        neg=df_Buffer_naNeg[df_Buffer_naNeg['buffer_conc']== buffer_conc]

        if len(pos['repeat'].unique())==3 and len(neg['repeat'].unique())==3:
            #calculate average no rna for buffer
            length=int(len(neg))
            arr = neg['value'].values.copy()
            arr.resize(3,int(length/3))
            DF_neg=pd.DataFrame(arr).T
            DF_neg['value_ave']=DF_neg.iloc[:, [0,1,2]].mean(axis=1)
            DF_neg['sem']=stats.sem(DF_neg.iloc[:, [0,1,2]].T)
            DF_neg['Time']=neg['Time'].reset_index(drop=True)[0:int(length/3)]*60

            #calculate average no rna for DTT
            length=int(len(neg))
            arr = pos['value'].values.copy()
            arr.resize(3,int(length/3))
            DF_pos=pd.DataFrame(arr).T
            DF_pos['value_ave']=DF_pos.iloc[:, [0,1,2]].mean(axis=1)-DF_neg['value_ave']
            DF_pos['sem']=stats.sem(DF_pos.iloc[:, [0,1,2]].T)
            DF_pos['Time']=pos['Time'].reset_index(drop=True)[0:int(length/3)]*3600

            # Add error bars to the DataFrame
            DF_pos['error_low'] = calibrateBiotek4(DF_pos['value_ave'] - DF_pos['sem'])
            DF_pos['error_high'] = calibrateBiotek4(DF_pos['value_ave'] + DF_pos['sem'])

            #Data from experiments
            pBuffer.line(x=DF_pos['Time']/3600,y=calibrateBiotek4(DF_pos['value_ave']),
            legend_label= (str(buffer_conc)), color= colorBlue[i], line_width=2,)
            # Add error bars
            pBuffer.segment(
                x0=DF_pos['Time']/3600, y0=DF_pos['error_low'],
                x1=DF_pos['Time']/3600, y1=DF_pos['error_high'],
                line_width=2, color= colorBlue[i],line_alpha=0.25,)
            i=i+1 
            
            mean=calibrateBiotek4(DF_pos['value_ave'][DF_pos['Time']<=6*3600].mean())
            sem=stats.sem(calibrateBiotek4(DF_pos['value_ave'][DF_pos['Time']<=6*3600]))
        elif buffer_conc=='0 mM':
            #calculate average no rna for Buffer
            length=int(len(neg))
            arr = neg['value'].values.copy()
            arr.resize(3,int(length/3))
            DF_neg=pd.DataFrame(arr).T
            DF_neg['value_ave']=DF_neg.iloc[:, [0,1,2]].mean(axis=1)
            DF_neg['sem']=stats.sem(DF_neg.iloc[:, [0,1,2]].T)
            DF_neg['Time']=neg['Time'].reset_index(drop=True)[0:int(length/3)]*60

            #calculate average no rna for Buffer
            length=int(len(neg))
            arr = pos['value'].values.copy()
            arr.resize(2,int(length/2))
            DF_pos=pd.DataFrame(arr).T
            DF_pos['value_ave']=DF_pos.iloc[:, [0,1]].mean(axis=1)-DF_neg['value_ave']
            DF_pos['sem']=stats.sem(DF_pos.iloc[:, [0,1]].T)
            DF_pos['Time']=pos['Time'].reset_index(drop=True)[0:int(length/2)]*3600

            # Add error bars to the DataFrame
            DF_pos['error_low'] = calibrateBiotek4(DF_pos['value_ave'] - DF_pos['sem'])
            DF_pos['error_high'] = calibrateBiotek4(DF_pos['value_ave'] + DF_pos['sem'])

            #Data from experiments
            pBuffer.line(x=DF_pos['Time']/3600,y=calibrateBiotek4(DF_pos['value_ave']),
            legend_label= (str(buffer_conc)+'*'), color= colorBlue[i], line_width=2,)
            # Add error bars
            pBuffer.segment(
                x0=DF_pos['Time']/3600, y0=DF_pos['error_low'],
                x1=DF_pos['Time']/3600, y1=DF_pos['error_high'],
                line_width=2, color= colorBlue[i], line_alpha=.25)
            i=i+1
            
            mean=calibrateBiotek4(DF_pos['value_ave'][DF_pos['Time']<=6*3600].mean())
            sem=stats.sem(calibrateBiotek4(DF_pos['value_ave'][DF_pos['Time']<=6*3600]))
            
        buffer_mean.append(mean)
        buffer_sem.append(sem)
    
    pBuffer.legend.location="top_right"
    pBuffer.legend.click_policy="hide"
    pBuffer.legend.border_line_color = None
    
    # print(buffer_mean)
    # bokeh.io.show(pBuffer)
    return ([pBuffer, buffer_mean, buffer_sem])
```

##### Get Data¶

In [19]:

```
# directory = r'C:/Users/zoila/Box/biocircuits/ZJurado/Projects/Organelles/PUREfrex_system/2023.07.23_3xBufferTest_redue_all/'
folder= '/2023.07.23_BufferswMGapt/'
filename = '/2023.07.23_BufferswMGapt_nPicklist.xlsx'
data_dict = pd.read_excel(directory + folder+filename, sheet_name=None, engine='openpyxl')
mgapt = data_dict['mgapt_tidy']
mgapt=mgapt[mgapt['comment']!= 'no mrna']

idxNegCon=mgapt['buffer_conc']== '0 mM'
```

##### Index and plot¶

In [20]:

```
#MGapt measurement over time in different buffers
plot=[]
buffer_Ave=pd.DataFrame({'conc': ['0 mM','1 mM','2 mM','4 mM',]})
buffer_Sem=pd.DataFrame({'conc': ['0 mM','1 mM','2 mM','4 mM',]})
for buffer in mgapt['buffer'].unique():
    if buffer != 'Neg' and buffer != 'PO4':
        p,ave,sem=BufferPlots(buffer, mgapt,idxNegCon)
        plot.append(p)
        buffer_Ave[buffer]=ave
        buffer_Sem[buffer]=sem
```

##### Plot as Grid and save as .svg¶

In [21]:

```
# using grid layout on plots
pBuffers=gridplot([[plot[0],plot[1],],[plot[2], plot[3],]], width=500, height=400)

count=1
for i in range(len(plot)):
    pic=plot[i]
    pic.output_backend = "svg"
    export_svgs(pic, filename = 'Figures_DTTvMGapt/Sup_pBuffers' + str(count) + '.svg',width=500, height=400)
    count+=1
export_svgs(pBuffers, filename = 'Figures_DTTvMGapt/Sup_pBuffers.svg')
show(pBuffers)
```

#### **Bar plot**¶

##### Definitions¶

In [22]:

```
def tosourcedata(df):
    buffers=['DTT','GSH','Ppi','TCEP']
    concs=['0 mM','1 mM','2 mM','4 mM',]
    sourcedata_df = {}
    sourcedata_df['buffer']= ['DTT','GSH','Ppi','TCEP']
    
    # Iterate over each concentration value
    for conc in df['conc']:
        # Select the row corresponding to the current concentration
        row = df[df['conc'] == conc][['DTT', 'GSH', 'Ppi', 'TCEP']].values.tolist()[0]
        # Store the sourcedata for the current concentration in the dictionary
        sourcedata_df[conc] = row
    return sourcedata_df
```

##### Initialize and plot¶

In [23]:

```
# # Initialize an empty dictionary to store sourcedata for each concentration
buffers=['DTT','GSH','PPi','TCEP']
concs=['0 mM','1 mM','2 mM','4 mM',]

data=tosourcedata(buffer_Ave)
error=tosourcedata(buffer_Sem)
```

In [24]:

```
x = [ (buffer, conc) for buffer in buffers for conc in concs ]
counts = sum(zip(data['0 mM'], data['1 mM'], data['2 mM'], data['4 mM']), ()) # like an hstack
error_counts = sum(zip(error['0 mM'], error['1 mM'], error['2 mM'], error['4 mM']), ()) # like an hstack
upper = [x+e for x,e in zip(counts, error_counts) ]
lower = [x-e for x,e in zip(counts, error_counts) ]

source = ColumnDataSource(data=dict(x=x, counts=counts, upper=upper, lower=lower))

p = figure(x_range=FactorRange(*x), height=350,#title="different buffers",
           toolbar_location=None, tools="",x_axis_label='Buffering Agents', y_axis_label='MGapt (μM)')

p.vbar(x='x', top='upper', width=0.05, source=source, line_color=None, fill_color='black')
p.vbar(x='x', top='lower', width=0.5, source=source, line_color=None, fill_color=factor_cmap('x', palette=colorBlue, factors=concs, start=1, end=2))
p.vbar(x='x', top='counts', width=0.9, source=source, line_color=None, fill_color=factor_cmap('x', palette=colorBlue, factors=concs, start=1, end=2))

p.y_range.start = 0
p.x_range.range_padding = 0.1
p.xaxis.major_label_orientation = 1
```

##### Save as .svg¶

In [25]:

```
# Set the output backend to SVG
p.output_backend = "svg"
p.ygrid.grid_line_color = None
p.xgrid.grid_line_color = None
p.outline_line_color = None

# Customize the font and size of the x-axis label
# p.xaxis.axis_label_text_font = "Arial"
p.xaxis.axis_label_text_font_size = "12pt"
p.xaxis.axis_label_text_font_style ='normal'
p.xaxis.axis_label_standoff = 10
# Customize the font and size of the y-axis label
# p.yaxis.axis_label_text_font = "Arial"
p.yaxis.axis_label_text_font_size = "12pt"
p.yaxis.axis_label_text_font_style ='normal'
p.yaxis.axis_label_standoff = 10

# Export the VBar plot as an SVG file
export_svgs(p, filename="Figures_DTTvMGapt/Fig_pBuffersAll.svg")
show(p)
```

### **Figure 3: Measure MGapt concentration dynamics**¶

#### a) Measure MGapt with known mRNA concentration¶

##### Get data¶

In [26]:

```
folder= '/2024.01.18_PURExpress_DTTconcs/'
filename = '2024.01.18_PURExpress_DDT.xlsx'
data_dict = pd.read_excel(directory + folder+ filename, sheet_name=None, engine='openpyxl')
sheets_tidy= [x for x in data_dict.keys() if '_tidy' in x]
df_mgapt=pd.read_csv(directory +  folder+'/'+ sheets_tidy[1]+'.csv')
```

```
C:\Users\zoila\anaconda3\envs\python38\lib\site-packages\openpyxl\worksheet\_reader.py:329: UserWarning: Unknown extension is not supported and will be removed
  warn(msg)
C:\Users\zoila\anaconda3\envs\python38\lib\site-packages\openpyxl\worksheet\_reader.py:329: UserWarning: Unknown extension is not supported and will be removed
  warn(msg)
```

##### Index and plot¶

In [27]:

```
neg=df_mgapt[df_mgapt['construct']=='none']
dna=df_mgapt[df_mgapt['na']=='dna']
rna=df_mgapt[df_mgapt['na']=='rna']

DF_Neg=Cal_avesNsems(neg)
```

In [28]:

```
# A:MGapt w/RNA
pRNA=create_custom_plot('a', x_max=12, y_max=0.5)

data=rna[rna['dtt_added']==0]
for na in data['construct'].unique():
    data2=data[data['construct']==na]
    DF=Cal_avesNsems(data2,DF_neg=DF_Neg)
    Circle_wErrorPlot(pRNA, DF, na, colorNA[na], cal=True, marker=markers[na],size=size[markers[na]])
# pRNA=updateLegend(pRNA,title='DNA construct:')
pRNA.legend.visible = False   
bokeh.io.show(pRNA)
```

##### Normalize¶

In [29]:

```
#B) MGapt w/RNA Norm
pRNAN=create_custom_plot('a', x_max=12, y_max=1,yname='Normalized MGapt')
data=rna[rna['dtt_added']==0]
for na in data['construct'].unique():
    data2=data[data['construct']==na]
    DF=Cal_avesNsems(data2,norm=True)
    Circle_wErrorPlot(pRNAN, DF,  na, colorNA[na], cal=False, marker=markers[na],size=size[markers[na]])
pRNAN.legend.visible = False
bokeh.io.show(pRNAN)
```

##### Save Legend¶

In [30]:

```
pConstruct=create_custom_plot('', x_max=12, y_max=1)
data=rna[rna['dtt_added']==0]
for na in data['construct'].unique():
    data2=data[data['construct']==na]
    DF=Cal_avesNsems(data2,norm=True)
    Circle_wErrorPlot(pConstruct, DF,  na, 'grey', cal=False, marker=markers[na],size=size[markers[na]])
# Get the legend from the plot
legend = pConstruct.legend[0]
legend.orientation = 'horizontal'
legend.location = 'center'
pConstruct.add_layout(legend, 'above')

for pic in [pConstruct]:
    pic.output_backend = "svg"
    export_svgs(pic, filename = 'Figures_DTTvMGapt/Fig3_pConstruct.svg',width=500, height=400)
export_svgs(pConstruct, filename = 'Figures_DTTvMGapt/Fig3_pConstruct.svg')
bokeh.io.show(pConstruct)
```

#### b) Experiment varying DNA concentration¶

##### Get data¶

In [31]:

```
folder= '/2024.01.26_PURExpress_DNAconcs/'
filename = '2024.01.26_PURExpress_DNAconcs.xlsx'
data_dict = pd.read_excel(directory + folder+filename, sheet_name=None, engine='openpyxl')
sheets_tidy= [x for x in data_dict.keys() if '_tidy' in x]
df_mgapt=pd.read_csv(directory + folder+'/'+ sheets_tidy[1]+'.csv')
df_mgapt=df_mgapt[df_mgapt['na_conc']!='na']
```

```
C:\Users\zoila\anaconda3\envs\python38\lib\site-packages\openpyxl\worksheet\_reader.py:329: UserWarning: Unknown extension is not supported and will be removed
  warn(msg)
C:\Users\zoila\anaconda3\envs\python38\lib\site-packages\openpyxl\worksheet\_reader.py:329: UserWarning: Unknown extension is not supported and will be removed
  warn(msg)
```

In [32]:

```
#C: Varying DNA
pDNAvar=create_custom_plot('b', x_max=12, y_max=3)
count=0
for na_conc in df_mgapt['na_conc'].unique():
    data=df_mgapt[df_mgapt['na_conc']==na_conc]
    na=data['construct'].unique()[0]
    DF=Cal_avesNsems(data,DF_neg=DF_Neg)
    Circle_wErrorPlot(pDNAvar, DF, str(na_conc)+' nM', colorPurples[5-count],cal=True,marker=markers[na],size=size[markers[na]])
    count=count+1
pDNAvar=updateLegend(pDNAvar,title='DNA (nM):')
bokeh.io.show(pDNAvar)
```

##### Normalize¶

In [33]:

```
#D: Varying DNA Normalized
pDNAvarN=create_custom_plot('b', x_max=12, y_max=1,yname='Normalized MGapt')
count=0
for na_conc in df_mgapt['na_conc'].unique():
    data=df_mgapt[df_mgapt['na_conc']==na_conc]
    na=data['construct'].unique()[0]
    DF=Cal_avesNsems(data,norm=True)
    Circle_wErrorPlot(pDNAvarN, DF, str(na_conc)+' nM', colorPurples[5-count],marker=markers[na],size=size[markers[na]])
    count=count+1
    
pDNAvarN=updateLegend(pDNAvarN,title='DNA (nM):')
bokeh.io.show(pDNAvarN)
```

#### c) Experiment varying DTT concentration with DNA¶

##### Get data¶

In [34]:

```
folder= '/2024.01.18_PURExpress_DTTconcs/'
filename = '2024.01.18_PURExpress_DDT.xlsx'
data_dict = pd.read_excel(directory + folder+ filename, sheet_name=None, engine='openpyxl')
sheets_tidy= [x for x in data_dict.keys() if '_tidy' in x]
df_mgapt=pd.read_csv(directory +  folder+'/'+ sheets_tidy[1]+'.csv')
```

```
C:\Users\zoila\anaconda3\envs\python38\lib\site-packages\openpyxl\worksheet\_reader.py:329: UserWarning: Unknown extension is not supported and will be removed
  warn(msg)
C:\Users\zoila\anaconda3\envs\python38\lib\site-packages\openpyxl\worksheet\_reader.py:329: UserWarning: Unknown extension is not supported and will be removed
  warn(msg)
```

##### Index and plot¶

In [35]:

```
neg=df_mgapt[df_mgapt['construct']=='none']
dna=df_mgapt[df_mgapt['na']=='dna']
rna=df_mgapt[df_mgapt['na']=='rna']

DF_Neg=Cal_avesNsems(neg)
```

In [36]:

```
#E: Varying DTT: DNA
test=pd.DataFrame(columns=DF.columns)
pDTTvar_dna=create_custom_plot('c', x_max=12, y_max=3.5)
count=0
for dtt_conc in dna['dtt_added'].unique():
    data=dna[dna['dtt_added']==dtt_conc]
    for na in data['construct'].unique():
        data2=data[data['construct']==na]
        DF=Cal_avesNsems(data2,DF_neg=DF_Neg)
        Circle_wErrorPlot(pDTTvar_dna, DF, str(dtt_conc)+' mM', colorDTT[3-count], cal=True, marker=markers[na],size=size[markers[na]])
    count += 1
# pDTTvar_dna=updateLegend(pDTTvar_dna,title='Additonal DTT (mM):')
pDTTvar_dna.legend.visible = False
bokeh.io.show(pDTTvar_dna)
```

##### Normalize¶

In [37]:

```
#F: Varying DTT: DNA Norm
pDTTvarN_dna=create_custom_plot('c', x_max=12, y_max=1,yname='Normalized MGapt')

count=0
for dtt_conc in dna['dtt_added'].unique():
    data=dna[dna['dtt_added']==dtt_conc]
    for na in data['construct'].unique():
        data2=data[data['construct']==na]
        DF=Cal_avesNsems(data2,norm=True)
        Circle_wErrorPlot(pDTTvarN_dna, DF, str(dtt_conc)+' mM', colorDTT[3-count], marker=markers[na],size=size[markers[na]])
    count += 1
pDTTvarN_dna.legend.visible = False    
bokeh.io.show(pDTTvarN_dna)
```

##### Save Legend¶

In [38]:

```
pDTTlabel=create_custom_plot('', x_max=12, y_max=1)
count=0
for dtt_conc in dna['dtt_added'].unique():
    data=dna[dna['dtt_added']==dtt_conc]
    for na in data['construct'].unique():
        data2=data[data['construct']==na]
        DF=Cal_avesNsems(data2,norm=True)
        Circle_wErrorPlot(pDTTlabel, DF, str(dtt_conc)+' mM', colorDTT[3-count], marker=markers[na],size=size[markers[na]])
    count += 1
# Get the legend from the plot
legend = pDTTlabel.legend[0]
legend.orientation = 'horizontal'
legend.location = 'center'
legend.title = "Additonal DTT (mM):"
legend.title_text_font_style = "normal"
pDTTlabel.add_layout(legend, 'above')

for pic in [pDTTlabel]:
    pic.output_backend = "svg"
    export_svgs(pic, filename = 'Figures_DTTvMGapt/Fig3_pDTTlabel.svg',width=500, height=400)
export_svgs(pDTTlabel, filename = 'Figures_DTTvMGapt/Fig3_pDTTlabel.svg')
bokeh.io.show(pDTTlabel)
```

#### d) Experiment varying DTT concentration with mRNA¶

In [39]:

```
#G: Varying DTT: RNA
pDTTvar_rna=create_custom_plot('d', x_max=12, y_max=0.5)

count=0
for dtt_conc in rna['dtt_added'].unique():
    data=rna[rna['dtt_added']==dtt_conc]
    for na in data['construct'].unique():
        data2=data[data['construct']==na]
        DF=Cal_avesNsems(data2,DF_neg=DF_Neg)
        Circle_wErrorPlot(pDTTvar_rna, DF, str(dtt_conc)+' mM', colorDTT[3-count], cal=True, marker=markers[na],size=size[markers[na]])
    count += 1
# pDTTvar_rna=updateLegend(pDTTvar_rna,title='Additonal DTT (mM):')
pDTTvar_rna.legend.visible = False  
bokeh.io.show(pDTTvar_rna)
```

##### Normalize¶

In [40]:

```
#H: Varying DTT: RNA Norm
pDTTvarN_rna=create_custom_plot('d', x_max=12, y_max=1,yname='Normalized MGapt')

count=0
for dtt_conc in rna['dtt_added'].unique():
    data=rna[rna['dtt_added']==dtt_conc]
    for na in data['construct'].unique():
        data2=data[data['construct']==na]
        DF=Cal_avesNsems(data2,norm=True)
        Circle_wErrorPlot(pDTTvarN_rna, DF, str(dtt_conc)+' mM', colorDTT[3-count], marker=markers[na],size=size[markers[na]])
    count += 1
pDTTvarN_rna.legend.visible = False   
bokeh.io.show(pDTTvarN_rna)
```

#### Plot as Grid and save as .svg¶

In [41]:

```
# using grid layout on plots
pMGaptExperiments=gridplot([[pRNAN, pDNAvarN,],[pDTTvarN_dna, pDTTvarN_rna]], width=500, height=400)

count=1
for pic in [pRNAN, pDNAvarN,  pDTTvarN_dna,pDTTvarN_rna]:
    pic.output_backend = "svg"
    export_svgs(pic, filename = 'Figures_DTTvMGapt/Fig3_pMGaptExp' + str(count) + '.svg',width=500, height=400)
    count+=1
export_svgs(pMGaptExperiments, filename = 'Figures_DTTvMGapt/Fig3_pMGaptExp.svg')
show(pMGaptExperiments)
```

In [42]:

```
# using grid layout on plots
pMGaptExperiments=gridplot([[pRNA,pDNAvar,],[pDTTvar_dna, pDTTvar_rna,]], width=500, height=400)

count=1
for pic in [pRNA, pRNAN, pDNAvar, pDNAvarN, pDTTvar_dna, pDTTvarN_dna,pDTTvar_rna,pDTTvarN_rna]:
    pic.output_backend = "svg"
    export_svgs(pic, filename = 'Figures_DTTvMGapt/Sup_pMGaptExp' + str(count) + '.svg',width=500, height=400)
    count+=1
export_svgs(pMGaptExperiments, filename = 'Figures_DTTvMGapt/Sup_pMGaptExp.svg')
show(pMGaptExperiments)
```

### **Figure 6: Model results overlayed with experiment results**¶

#### a,b) Simulation vs. experimental results¶

##### Get data¶

In [43]:

```
# Getting experimental data
folder= '/Exp_mRNA_concs/'
## Negative control
filename0 = 'Data_0.csv'
DF_Neg =  pd.read_csv(directory+ folder+  filename0, delimiter = '\,', names = ['time','mRNA0'], engine='python',skiprows = 1)
DF_Neg=DF_Neg.astype(float)
```

##### Get modeling¶

In [44]:

```
#Upload model
modelname="Model_DTTvMGapt_Final"
M_fit = Model(sbml_filename =  modelname+"_updated.xml")

# Custom plots using mcmc_results.csv
labels = ['$k_{MGapt}$','$kb_{DTT}$',
          '$ku_{mgapt}$','$kb_{mgapt}$','$k_{DTTdeg}$',]
samples_mcmc = pd.read_csv('mcmc_results_Final.csv', names = labels,
                           engine = 'python')
#Sample over distribution
samples_discard = samples_mcmc.tail(500) #burning samples; take last 500
samples_mean = samples_discard.mean() #taking mean of the samples

for pi, pi_val in zip(['k_forward','k_forward1',
                           'k_forward3','k_reverse4','k_forward6',], samples_mean):
    M_fit.set_parameter(pi, pi_val)

# Run model to test
# p1=create_custom_plot('MGapt fit', x_max=8, y_max=1.6)
timepoints = np.linspace(0, 12*3600, 1200)
multMGapt=0.45 #From previous workbook
```

##### Plot data over model results¶

In [45]:

```
#Model at different initial mRNA concentrations
p1=create_custom_plot('a', x_max=12, y_max=1.6,yname='MGapt (measured, μM)')
p2=create_custom_plot('b', x_max=4, y_max=1.6, yname='MGapt (measured, μM)')
na_conc=[0.22, 0.41, 0.86, 1.26, 1.67]

for i in range(len(na_conc)):
    filename = 'Data_'+str(na_conc[i])+'.csv'
    mgapt =  pd.read_csv(directory+ folder+ filename, delimiter = '\,', names = ['time','mRNA0', 'mRNA1','mRNA2',], engine='python',skiprows = 1)
    mgapt=mgapt.astype(float)[:145]
    
    DF_MGapt=Cal_avesNsems2(mgapt,DF_neg=DF_Neg)
    Circle_wErrorPlot(p1, DF_MGapt, str(na_conc[i])+' μM', colorsmRNA[i],cal=True)
    Circle_wErrorPlot(p2, DF_MGapt, str(na_conc[i])+' μM', colorsmRNA[i],cal=True)
    
    #Run Model
    initial_con={'MGapt':(na_conc[i]*multMGapt), 'MGapt_altered':(na_conc[i]*(1-multMGapt)),'DTT':(1000),'MGdye':(10-na_conc[i])} #in uM
    M_fit.set_species(initial_con)
    Model_DTT = py_simulate_model(Model = M_fit, timepoints = timepoints) 
    Model_DTT['Measured']=Model_DTT['MGapt']+Model_DTT['MGapt_altered']*.15
    Model_DTT.to_csv(directory0+'/Simulation_Results/rnaMGapt-deGFP_'+str(na_conc[i])+'.csv') 
    p1.line(timepoints/3600, Model_DTT['Measured'], color= colorsmRNA[i], legend_label=str(na_conc[i])+' μM',)
    p2.line(timepoints/3600, Model_DTT['Measured'], color= colorsmRNA[i], legend_label=str(na_conc[i])+' μM',)

p1.legend.location="top_left"
p1.legend.click_policy="hide"
p1.legend.border_line_color = None
bokeh.io.show(p1)
```

##### Plot as Grid and save as .svg¶

In [46]:

```
# using grid layout on plots
pResults=gridplot([[p1, p2]], width=500, height=400)

count=1
for pic in [p1, p2]:
    pic.output_backend = "svg"
    export_svgs(pic, filename = 'Figures_DTTvMGapt/Fig6_pResults' + str(count) + '.svg',width=500, height=400)
    count+=1
export_svgs(pResults, filename = 'Figures_DTTvMGapt/Fig6_pResults.svg')
show(pResults)
```

#### Plot Errors¶

In [47]:

```
# Absolute and percent Error
pAbsError=create_custom_plot('a', x_max=4, y_max=.14, yname='Difference in Measured MGapt (μM)')
pError=create_custom_plot('b', x_max=4, y_max=50, yname='Error (%)')

timepoints = np.array(DF_MGapt['Time'])

for i in range(len(na_conc)):
    #Get data
    filename = 'Data_'+str(na_conc[i])+'.csv'
    mgapt =  pd.read_csv(directory+ folder+ filename, delimiter = '\,', names = ['time','mRNA0', 'mRNA1','mRNA2',], engine='python',skiprows = 1)
    mgapt=mgapt.astype(float)[:145]
    DF_MGapt=Cal_avesNsems2(mgapt,DF_neg=DF_Neg)
    
    #Get Model
    initial_con={'MGapt':(na_conc[i]*multMGapt), 'MGapt_altered':(na_conc[i]*(1-multMGapt)),'DTT':(1000),'MGdye':(10-na_conc[i])} #in uM
    M_fit.set_species(initial_con)
    Model_DTT = py_simulate_model(Model = M_fit, timepoints = timepoints) 
    Model_DTT['Measured']=Model_DTT['MGapt']+Model_DTT['MGapt_altered']*.15
    
    #Calculate Error
    DF_MGapt['uM_ave']=calibrateBiotek4(DF_MGapt['value_ave'])
    diff_mgapt=np.abs(DF_MGapt['uM_ave']-Model_DTT['Measured'])
    pAbsError.scatter(timepoints/3600, diff_mgapt, color= colorsmRNA[i], legend_label=str(na_conc[i])+' μM',fill_alpha=.5)
    pAbsError.line(timepoints/3600, diff_mgapt, color= colorsmRNA[i], legend_label=str(na_conc[i])+' μM',line_alpha=.5,line_dash='dashed', )    
    
    percent_error=diff_mgapt/DF_MGapt['uM_ave']*100
    print(f'Average Percent Error of {str(na_conc[i])} μM: {percent_error.mean()}')
    pError.scatter(timepoints/3600, percent_error, color= colorsmRNA[i], legend_label=str(na_conc[i])+' μM',fill_alpha=.5)
    pError.line(timepoints/3600, percent_error, color= colorsmRNA[i], legend_label=str(na_conc[i])+' μM',line_alpha=.5,line_dash='dashed', )    
    
pAbsError.legend.location="top_left"
pAbsError.legend.click_policy="hide"
pAbsError.legend.border_line_color = None

pError.legend.location="top_left"
pError.legend.click_policy="hide"
pError.legend.border_line_color = None
```

```
Average Percent Error of 0.22 μM: 8.268444299994057
Average Percent Error of 0.41 μM: 6.159201825832312
Average Percent Error of 0.86 μM: 5.775026250384164
Average Percent Error of 1.26 μM: 6.972065889657352
Average Percent Error of 1.67 μM: 9.752354336510683
```

##### Plot as Grid and save as .svg¶

In [48]:

```
# using grid layout on plots
pErrorAll=gridplot([[pAbsError, pError]], width=500, height=400)

count=1
for pic in [pAbsError, pError]:
    pic.output_backend = "svg"
    export_svgs(pic, filename = 'Figures_DTTvMGapt/Sup_pError' + str(count) + '.svg',width=500, height=400)
    count+=1
export_svgs(pResults, filename = 'Figures_DTTvMGapt/Sup_pError.svg')
show(pErrorAll)
```

### **Back Calculate**¶

In [49]:

```
# Run model to test
# A) Experimental and modeled data of Measured MGapt
pRNA22=create_custom_plot('a', x_max=4, y_max=.25,)
# B) Dynamic Calibration Curve of MGapt
pRatio=create_custom_plot('b', x_max=4, y_max=8, yname='Ratio of MGapt (Input/Modeled)')
# C) Experimental Data of Measured MGapt'
pData=create_custom_plot('c', x_max=4, y_max=2,)
# D) Dynamic Calibration of Experimental Data
pData_Cal=create_custom_plot('d', x_max=4, y_max=2,)
timepoints = np.array(DF_MGapt['Time'])

for i in range(len(na_conc)):
    #Get data
    filename = 'Data_'+str(na_conc[i])+'.csv'
    mgapt =  pd.read_csv(directory+ folder+ filename, delimiter = '\,', names = ['time','mRNA0', 'mRNA1','mRNA2',], engine='python',skiprows = 1)
    mgapt=mgapt.astype(float)[:145]
    DF_MGapt=Cal_avesNsems2(mgapt,DF_neg=DF_Neg)
    
    #Get Model
    initial_con={'MGapt':(na_conc[i]*multMGapt), 'MGapt_altered':(na_conc[i]*(1-multMGapt)),'DTT':(1000),'MGdye':(10-na_conc[i])} #in uM
    M_fit.set_species(initial_con)
    Model_DTT = py_simulate_model(Model = M_fit, timepoints = timepoints) 
    Model_DTT['Measured']=Model_DTT['MGapt']+Model_DTT['MGapt_altered']*.15
    
    #Calculate Calibration Curve
    if i==0:
        pRNA22.line(timepoints/3600, Model_DTT['Measured'], color= colorsmRNA[i], legend_label='Simulated data: '+str(na_conc[i])+' μM',  line_alpha=.5, line_width=2)
    
        cal_curve=na_conc[i]/Model_DTT['Measured']
        pRatio.square(timepoints/3600, cal_curve, color= colorsmRNA[i], legend_label='(Experimental data)/(Simulated data): '+str(na_conc[i])+' μM',  fill_alpha=0.5)
        
        Circle_wErrorPlot(pRNA22, DF_MGapt, 'Experimental data: '+str(na_conc[i])+' μM', colorsmRNA[i],cal=True)
    else:
        Circle_wErrorPlot(pData, DF_MGapt, str(na_conc[i])+' μM', colorsmRNA[i],cal=True)
        
    #Calculate "True" MGapt
    if i!=0:
        DF_MGapt_new=pd.DataFrame(columns=['Time','uM_ave', 'error_low', 'error_high'])
        DF_MGapt_new['Time']=DF_MGapt['Time']
        DF_MGapt_new['value_ave']=calibrateBiotek4(DF_MGapt['value_ave'])*cal_curve
        DF_MGapt_new['error_low']=calibrateBiotek4(DF_MGapt['error_low'])*cal_curve
        DF_MGapt_new['error_high']=calibrateBiotek4(DF_MGapt['error_high'])*cal_curve
        
        Circle_wErrorPlot(pData_Cal, DF_MGapt_new, str(na_conc[i])+' μM', colorsmRNA[i],cal=False)
        pData_Cal.line(timepoints/3600, na_conc[i], line_dash='dashed',  color= colorsmRNA[i], legend_label=str(na_conc[i])+' μM',line_alpha=.5)
        
        # Add text
        labels = Label(x=3.25, y=na_conc[i], text=str(na_conc[i])+' μM',)
        pData_Cal.add_layout(labels)

    
    
pRNA22.legend.location="top_right"
pRNA22.legend.click_policy="hide"
pRNA22.legend.border_line_color = None

pRatio.legend.location="top_right"
pRatio.legend.click_policy="hide"
pRatio.legend.border_line_color = None

pData.legend.location="top_right"
pData.legend.click_policy="hide"
pData.legend.border_line_color = None

# Remove the legend
pData_Cal.legend.visible = False
```

#### Plot as Grid and save as .svg¶

In [50]:

```
# using grid layout on plots
pBackCalc=gridplot([[pRNA22, pRatio],[pData, pData_Cal]], width=500, height=400)
count=1
for pic in [pRNA22, pRatio,pData, pData_Cal]:
    pic.output_backend = "svg"
    export_svgs(pic, filename = 'Figures_DTTvMGapt/Sup_pBackCalc' + str(count) + '.svg',width=500, height=400)
    count+=1
export_svgs(pBackCalc, filename = 'Figures_DTTvMGapt/Sup_pBackCalc.svg')
show(pBackCalc)
```

### **Calibrations Plots**¶

#### mRNA Calibrations¶

In [51]:

```
#MGapt Calibration file
filename = '/Calibrations_Biotek/2022.10.04_mRNA_mGapt_data_37C.csv'
mGapt_Data =  pd.read_csv(directory0+filename, delimiter = '\,', 
                          names = ['mRNA (nM)','Biotek 1', 'Biotek 2','Biotek 3', 'Biotek 4'], 
                          skiprows = 1,engine='python')
```

In [52]:

```
pCal_mRNA = bokeh.plotting.figure(toolbar_location='right',
    outline_line_color= None,
    min_border_right=10,
    height=400,
    width=500,)

# pCal_mRNA.title.text = "A) MGapt Calibration: Ex/Em 610/650, Gain 150 "
pCal_mRNA.xaxis.axis_label = 'mRNA (nM)'
pCal_mRNA.yaxis.axis_label = 'Fluorescence (a.u.)'
pCal_mRNA.y_range=Range1d(0, 1400)
pCal_mRNA.x_range=Range1d(0, 65)
pCal_mRNA.outline_line_color=None

# pCal_mRNAyaxis
pCal_mRNA.ygrid.visible = False
pCal_mRNA.yaxis.axis_label_text_font_size='15pt' 
pCal_mRNA.yaxis.major_label_text_font_size = '15pt'
pCal_mRNA.yaxis.major_label_text_font='Work Sans'
pCal_mRNA.yaxis.axis_label_standoff=15
pCal_mRNA.yaxis.axis_label_text_font_style='normal' 

# pCal_mRNAxaxis
pCal_mRNA.xgrid.visible = False
pCal_mRNA.xaxis.axis_label_text_font_size='15pt' 
pCal_mRNA.xaxis.major_label_text_font_size = '15pt'
pCal_mRNA.xaxis.major_label_text_font='Work Sans'
pCal_mRNA.xaxis.axis_label_standoff=15
pCal_mRNA.xaxis.axis_label_text_font_style='normal' 

# pCal_mRNAtitle
pCal_mRNA.title.text_font_size= '18pt' 
pCal_mRNA.title.align= 'left'
pCal_mRNA.title.offset=-70.0


#Data from experiments
pCal_mRNA.scatter(mGapt_Data['mRNA (nM)'], mGapt_Data['Biotek 4'], radius=0.5, fill_alpha=0.5, color="navy") #mRNA
m4, b4 = np.polyfit(mGapt_Data['mRNA (nM)'], mGapt_Data['Biotek 4'], 1)
pCal_mRNA.line(mGapt_Data['mRNA (nM)'],mGapt_Data['mRNA (nM)']*m4+b4 ,  color="#d95b43", line_dash= 'dashed', line_width=2) #trendline

citation4 = Label(x=200, y=100, x_units='screen', y_units='screen',
                 text='y= '+str(np.around(m4, decimals=2, out=None))+'x + ' + str(np.around(b4, decimals=2, out=None)),# render_mode='css',
                 border_line_color=None, border_line_alpha=1.0,
                 background_fill_color='white', background_fill_alpha=1.0, text_font_size='14pt')
pCal_mRNA.add_layout(citation4)

bokeh.io.show(pCal_mRNA)
```

In [53]:

```
count=1
for pic in [pCal_mRNA]:
    pic.output_backend = "svg"
    export_svgs(pic, filename = 'Figures_DTTvMGapt/Sup_pCalB4.svg',width=500, height=400)
    count+=1
export_svgs(pCal_mRNA, filename = 'Figures_DTTvMGapt/Sup_pCalB4.svg', width=500, height=400)
```

Out[53]:

```
['Figures_DTTvMGapt/Sup_pCalB4.svg']
```

### **SNARF Plots**¶

##### Get data¶

In [54]:

```
#SNARF experiment files
folder = '/SNARF-5F_PURExpress/'
df_gfp=pd.read_csv(directory0 +  folder+ 'gfp61_data_tidy.csv')
df_snarf580=pd.read_csv(directory0 +  folder+'SNARF580_data_tidy.csv')
df_snarf640=pd.read_csv(directory0 +  folder+'SNARF640_data_tidy.csv')
```

##### Index and plot¶

In [55]:

```
idxP=df_gfp['system']=='PURExpress'
idxS=df_gfp['system']=='snarf-5f'

df_snarf580cal=df_snarf580[idxS]
df_snarf640cal=df_snarf640[idxS]

df_snarf580data=df_snarf580[idxP]
df_snarf640data=df_snarf640[idxP]
```

#### Plots SNARF calibration over time¶

In [56]:

```
#SNARF-5F over time
pSNARF_cal=create_custom_plot('a', x_max=12, y_max=3.5,yname='RFU (580/640)')
i=0
for well in df_snarf_cal['well'].unique():
    idxw=df_snarf_cal['well']==well
    
    pSNARF_cal.scatter(df_snarf_cal[idxw]['time'], df_snarf_cal[idxw]['580/640'], legend_label =  df_snarf_cal[idxw]['pH'].unique()[0], 
              color=colorspH[i], alpha=.5)
    i+=1
# Get the legend from the plot
pSNARF_cal=updateLegend(pSNARF_cal,title='pH:',layout_loc='right')
bokeh.io.show(pSNARF_cal)
```

```
---------------------------------------------------------------------------
NameError                                 Traceback (most recent call last)
Cell In[56], line 4
      2 pSNARF_cal=create_custom_plot('a', x_max=12, y_max=3.5,yname='RFU (580/640)')
      3 i=0
----> 4 for well in df_snarf_cal['well'].unique():
      5     idxw=df_snarf_cal['well']==well
      7     pSNARF_cal.scatter(df_snarf_cal[idxw]['time'], df_snarf_cal[idxw]['580/640'], legend_label =  df_snarf_cal[idxw]['pH'].unique()[0], 
      8               color=colorspH[i], alpha=.5)

NameError: name 'df_snarf_cal' is not defined
```

#### Plots SNARF calibration¶

In [ ]:

```
#SNARF Calibration for pH
pSNARF_ave=create_custom_plot('b', x_max=10, y_max=3.5,xname= 'pH',yname='RFU (580/640)')

df_snarf=pd.DataFrame(columns=['pH','value_ave','sem','error_low','error_high'])
df_snarf['pH']=df_snarf_cal['pH'].unique()[:11]
i=0
for pH in df_snarf_cal['pH'].unique():
    df_cal=df_snarf_cal[df_snarf_cal['pH']==pH]
    df_snarf['value_ave'][i]=df_cal['580/640'].mean()
    df_snarf['sem'][i]=stats.sem(df_cal['580/640'])
    df_snarf['error_low'][i]=df_cal['580/640'].mean()-stats.sem(df_cal['580/640'])
    df_snarf['error_high'][i]=df_cal['580/640'].mean()+stats.sem(df_cal['580/640'])
    i+=1
# Plot
pSNARF_ave.scatter(
        x=df_snarf['pH'], y=(df_snarf['value_ave']), color='black', marker="circle",size=5, fill_alpha=0.2,)
# Add error bars
pSNARF_ave.segment(
    x0=df_snarf['pH'], y0=(df_snarf['error_low']),
    x1=df_snarf['pH'], y1=(df_snarf['error_high']),
    line_width=2, color= 'black', line_alpha=.25)

#add equation
start=2
end=5
y = df_snarf['value_ave'][start:end].astype(float)
x = df_snarf['pH'][start:end].astype(float)
m,b = np.polyfit(x, y, 1)
y2=m*x+b
pSNARF_ave.line(df_snarf['pH'][start:end], y2,line_dash='dashed', line_color='black') 
pSNARF_ave.x_range.start = 2.5
mytext = Label(x=7.5, y=.5, text='y='+str(np.round(m,2))+'x+'+str(np.round(b,2)),)
pSNARF_ave.add_layout(mytext)

bokeh.io.show(pSNARF_ave)
```

#### Plots Expression calibration over time¶

In [ ]:

```
df_snarf_data=pd.DataFrame() 
df_snarf_data['time']=df_snarf580data['Time']
df_snarf_data['well']=df_snarf580data['well']
df_snarf_data['construct']=df_snarf580data['construct']
df_snarf_data['580']=df_snarf580data['value']
df_snarf_data['640']=df_snarf640data['value']
df_snarf_data['580/640']=df_snarf_data['580']/df_snarf_data['640']
df_snarf_data['pH']= (df_snarf_data['580/640']-b)/m

pSNARF=create_custom_plot('c', x_max=12, y_max=8,yname='pH')

for well in df_snarf_data['well'].unique():
    idxw=df_snarf_data['well']==well
    if df_snarf_data[idxw]['construct'].unique()!='pT7-sfCFP':
        name=df_snarf_data[idxw]['construct'].unique()[0]
        pSNARF.scatter(df_snarf_data[idxw]['time'], df_snarf_data[idxw]['pH'], legend_label =  df_snarf_data[idxw]['construct'].unique()[0], 
                  color=colorSNARF[name], alpha=.5)
pSNARF.legend.title = 'Expressing from:'
pSNARF.legend.title_text_font_style = "normal"
pSNARF.y_range.start = 7.75 
pSNARF.legend.location="top_right"
pSNARF.legend.click_policy="hide"
pSNARF.legend.border_line_color = None
bokeh.io.show(pSNARF)
```

#### Plots deGFP during SNARF measurements¶

In [ ]:

```
#Get data for gfp measurements only with PURE
gfp=df_gfp[idxP]


pSNARF_gfp=create_custom_plot(' d', x_max=12, y_max=6,yname='deGFP (μM)')

for well in gfp['well'].unique():
    idxw=gfp['well']==well
    name=gfp[idxw]['construct'].unique()[0]
    if name!='pT7-sfCFP':
        pSNARF_gfp.scatter(gfp[idxw]['Time'], gfp[idxw]['value']/6176, 
                  color=colorSNARF[name], alpha=.5)
        # Add text
        labels = Label(x=8.5, y=(gfp[idxw]['value']/6176).max(), text=name,text_font_size='10pt')
        pSNARF_gfp.add_layout(labels)
        
bokeh.io.show(pSNARF_gfp)
```

#### Plot as Grid and save as .svg¶

In [ ]:

```
# using grid layout on plots
pSNARFS=gridplot([[pSNARF_cal, pSNARF_ave],[pSNARF, pSNARF_gfp]], width=500, height=400)
count=1
for pic in [pSNARF_cal, pSNARF_ave,pSNARF, pSNARF_gfp]:
    pic.output_backend = "svg"
    export_svgs(pic, filename = 'Figures_DTTvMGapt/Sup_pSNARFS' + str(count) + '.svg',width=500, height=400)
    count+=1
export_svgs(pSNARFS, filename = 'Figures_DTTvMGapt/Sup_pSNARFS.svg')
show(pSNARFS)
```

### Computing environment¶

In [ ]:

```
%load_ext watermark
%watermark -v -p bioscrape,bokeh,panel,jupyterlab,biocrnpyler
```
