## Supplementary Data Files for "Impact of Chemical Dynamics of Commercial PURE Systems on Malachite Green Aptamer Fluorescence": Model_DTTvMGapt_Sensivitiy_Inference_Final.html

March 26, 2024

[1] Division of Engineering and Applied Science, California Institute of Technology, Pasadena, CA \

In this notebook, we built a chemical reaction network for the interaction of DTT and MGapt.

### Importing required packages and definitions¶

In [1]:

```
#General
import corner
import matplotlib.pyplot as plt 
import numpy as np
import pandas as pd
import warnings
import seaborn as sn
import time
from scipy.integrate import odeint
import libsbml

#From BioCRNpyler
import biocrnpyler
from biocrnpyler import *
from biocrnpyler.component import Component
from biocrnpyler.chemical_reaction_network import Species, Reaction, ChemicalReactionNetwork
from biocrnpyler.mechanism import Mechanism
from biocrnpyler.reaction import Reaction
from biocrnpyler.species import Complex, Species, WeightedSpecies

#From bioscrape
from bioscrape.simulator import py_simulate_model, ModelCSimInterface, DeterministicSimulator
from bioscrape.inference import py_inference
from bioscrape.analysis import py_sensitivity_analysis
from bioscrape.types import Model

#Get directory
import os
directory = os.getcwd()
```

#### Bokeh¶

In [2]:

```
%matplotlib inline
try:
    import dnaplotlib as dpl
    dpl_enabled = True
except (ModuleNotFoundError,ImportError) as e:
    dpl_enabled = False
    
# Modules needed from Bokeh.
import bokeh.io
bokeh.io.output_notebook()   
import bokeh.plotting
from bokeh.themes import Theme
from bokeh.io import output_file, show
from bokeh.plotting import figure
from bokeh.models import LinearAxis, Range1d
from bokeh.io import export_png

#Colors
colors= {'mRNA0':"lightblue", 'mRNA1':"cornflowerblue",'mRNA2':"steelblue"}
colors2 = bokeh.palettes.Colorblind[8]
```

Loading BokehJS ...

#### Definitions¶

In [3]:

```
#Definition: Allows models built in BioCRNpyler to be rewritten with global reaction rates. 
## The updated .xml files can then be used in sensitivity anaylsis and inferencing. 
def update_model_local_to_global_params(filename):

    # rename parameters in sbml file so that they have same ID
    sbml_doc = libsbml.readSBMLFromFile(filename) # load your old SBML file
    param_dict = {}
    sbml_model = sbml_doc.getModel()
    for rxn in sbml_model.getListOfReactions():
        kl = rxn.getKineticLaw()
        for param in kl.getListOfLocalParameters():
            param_dict[param.getValue()] = param.getId()
    added_params = []
    changelog = {}
    count = 0
    for p_val, p in param_dict.items():
        sbml_param = sbml_model.createParameter()
        if p in added_params:
            p_new = p+str(count)
            changelog[p_val] = p_new
            sbml_param.setId(p_new)
            added_params.append(p_new)
        else:
            sbml_param.setId(p)
            added_params.append(p)
        count += 1
        sbml_param.setValue(p_val)
        sbml_param.setConstant(False)

    for rxn in sbml_model.getListOfReactions():
        kl = rxn.getKineticLaw()
        for param in list(kl.getListOfLocalParameters()):
            if param.getValue() in changelog:
                # print('renaming {0} with {1}'.format(param.getId(), changelog[param.getValue()]))
                kl.setFormula(kl.getFormula().replace(param.getId(), changelog[param.getValue()]))
            kl.removeLocalParameter(param.getId())


    updated_filname = filename.split('.xml')[0] + '_updated.xml'

    libsbml.writeSBML(sbml_doc, updated_filname) # write your new SBML
```

In [4]:

```
def create_custom_plot(title_text, x_max=8,y_max=2, yname='MGapt (μM)'):
    custom_plot = figure(
        toolbar_location='right',
        outline_line_color=None,
        min_border_right=10,
        height=400,
        width=500,)

    custom_plot.title.text = title_text
    custom_plot.xaxis.axis_label = 'Time (hours)'
    custom_plot.yaxis.axis_label = yname
    custom_plot.y_range = Range1d(0, y_max)
    custom_plot.x_range = Range1d(0, x_max)
    custom_plot.outline_line_color = None

    # custom_plot.yaxis
    custom_plot.ygrid.visible = False
    custom_plot.yaxis.axis_label_text_font_size = '15pt'
    custom_plot.yaxis.major_label_text_font_size = '15pt'
    custom_plot.yaxis.major_label_text_font = 'Work Sans'
    custom_plot.yaxis.axis_label_standoff = 15
    custom_plot.yaxis.axis_label_text_font_style = 'normal'

    # custom_plot.xaxis
    custom_plot.xgrid.visible = False
    custom_plot.xaxis.axis_label_text_font_size = '15pt'
    custom_plot.xaxis.major_label_text_font_size = '15pt'
    custom_plot.xaxis.major_label_text_font = 'Work Sans'
    custom_plot.xaxis.axis_label_standoff = 15
    custom_plot.xaxis.axis_label_text_font_style = 'normal'

    # custom_plot.title
    custom_plot.title.text_font_size = '18pt'
    custom_plot.title.align = 'left'
    custom_plot.title.offset = -70.0

    return custom_plot
```

### 1. CRN model of DTT and MGapt interaction¶

In [5]:

```
#Species
mgapt=Species('mgapt') # Malachite green aptamer 
MGapt=Species('MGapt') # Malachite green aptamer bound to MG-dye
DTT=Species('DTT')# Dithiothreitol
MGapt_altered=Species('MGapt_altered')# Altered Malachite green aptamer (less fluorescent)
MGdye=Species('MGdye')# Malachite green

species_DTTvMGapt=[MGapt, DTT, MGapt_altered,mgapt, MGdye]

# Reaction rates
k_DTTdeg = 3.2e-4     # (k_forward)  degradation rate of DTT
kb_DTT   = 1.935e-06  # (k_forward1) binding of DTT to MGapt
ku_DTT   = 7.74e-06   # (k_reverse)  unbinding of DTT to MGapt
k_MGapt  = 1.0e-3     # (k_forward3) MG-dye binding to mRNA
ku_mgapt = 0.0774     # (k_forward4) unbinding of DTT from altered Malachite green aptamer
kb_mgapt = 0.07740774 # (k_reverse5) rebinding of DTT to make the altered Malachite green aptamer
k_RNAdeg = 20.0e-4   # (k_forward6) degradation rate of unfolded mRNA

rxns_DTTvMGapt=[
        Reaction.from_massaction([mgapt,MGdye], [MGapt], k_forward=k_MGapt,),
        Reaction.from_massaction([MGapt,DTT], [MGapt_altered], k_forward=kb_DTT, k_reverse=ku_DTT),
        Reaction.from_massaction([MGapt_altered], [mgapt, DTT,MGdye], k_forward=ku_mgapt,k_reverse=kb_mgapt),
        Reaction.from_massaction([mgapt], [], k_forward=k_RNAdeg),
        Reaction.from_massaction([DTT], [], k_forward=k_DTTdeg),] 

#Compile model
CRN_DTTvMGapt = ChemicalReactionNetwork(species = species_DTTvMGapt, reactions = rxns_DTTvMGapt)
#Save model
filename="Model_DTTvMGapt_Final"
CRN_DTTvMGapt.write_sbml_file(filename+".xml")
update_model_local_to_global_params(filename+".xml")
```

#### Simulate model and plot¶

In [6]:

```
# Run model
start_time = time.time() #Start time
timepoints = np.linspace(0, 8*3600, 800)
mRNA_total=.22 #total mRNA added by user or in reaction
initial_con={'MGapt':(mRNA_total*.5), 'MGapt_altered':(mRNA_total*.5),'DTT':(1000),'MGdye':(10-mRNA_total)} #in uM
Results_DTTvMGapt = CRN_DTTvMGapt.simulate_with_bioscrape_via_sbml(timepoints = timepoints, initial_condition_dict = initial_con)

end_time = time.time()  #Endtime
total_time = end_time - start_time #Elapsed time
print(f"Simulation Time: {total_time} seconds")

# Calculate the simulated measurement of fluorescence
Results_DTTvMGapt['Measured']=Results_DTTvMGapt['MGapt']+Results_DTTvMGapt['MGapt_altered']*.15

#Setup plot
pMGapt=create_custom_plot('MGapt measured by Biotek', x_max=8, y_max=0.25)
pMGapt.line(timepoints/3600, Results_DTTvMGapt['Measured'])
bokeh.io.show(pMGapt)
```

```
Simulation Time: 0.03167152404785156 seconds
```

### 2. Sensitivity Analysis¶

*add some words*

In [7]:

```
#Get model
m = Model(sbml_filename = filename+"_updated.xml")
m.set_species(initial_con)

#Run sensitivity analysis
timepoints = np.linspace(0, 8*3600, 800)
ssm = py_sensitivity_analysis(model = m, timepoints = timepoints, normalize = True)
SSM=np.around(ssm[:,:,2].T, decimals=3, out=None)
```

```
C:\Users\zoila\anaconda3\envs\python38\lib\site-packages\bioscrape\analysis.py:244: RuntimeWarning: invalid value encountered in divide
  SSM_normalized[:,j,i] = np.divide(SSM[:,j,i]*params_values[j], solutions[:,i])
```

In [21]:

```
#Plot 
figsize = (15,8)
fig, ax = plt.subplots(figsize=figsize)
index = 1
params_names_latex = ['$k_{MGapt}$','$kb_{DTT}$','$ku_{DTT}$',
                      '$ku_{mgapt}$','$kb_{mgapt}$','$k_{RNAdeg}$','$k_{DTTdeg}$',]

# Round the timepoints
rounded_timepoints = np.round(np.linspace(timepoints[0] / 3600, timepoints[-1] / 3600, len(timepoints),
                                           endpoint=True, dtype='float'), 1)

sn_ax = sn.heatmap(SSM, ax = ax, annot=False, cmap = 'YlGnBu',
                   xticklabels = rounded_timepoints, center=0)

# title = sn_ax.set_title("a", fontsize=15, loc='left')
title = ax.set_title(
    "a", loc="left", y=.98,x=-.05,
    rotation=0, ha="left", va="center",fontsize=18,)

sn_ax.figure.axes[-1].xaxis.label.set_size(20)
ax = fig.axes
_ = plt.xlabel('Time (h)', fontsize = 15)
_ = plt.ylabel('Parameters', fontsize = 15)
_ = ax[0].tick_params(axis='x', which='major', labelsize=15, bottom = False)
_ = ax[0].tick_params(axis='y', which='major', labelsize=15, left = False)
_ = ax[0].set_yticklabels(params_names_latex)
every_nth = 50
for n, label in enumerate(ax[0].xaxis.get_ticklabels()):
    if n == len(timepoints)-1:
        continue
    if n % every_nth != 0:
        label.set_visible(False)
_ = ax[1].tick_params(axis = 'x', labelsize = 18)
_ = plt.savefig('Figures_DTTvMGapt/MGaptvDTT-sensitivity.svg')
plt.show()
```

In [9]:

```
for x in range(len(params_names_latex)):
    print(params_names_latex[x], SSM[x,1:].mean())
```

```
$k_{MGapt}$ -0.44825281602002504
$kb_{DTT}$ 0.16183103879849817
$ku_{DTT}$ -0.02056821026282854
$ku_{mgapt}$ -0.5634555694618273
$kb_{mgapt}$ 0.5491927409261577
$k_{RNAdeg}$ -0.10012015018773467
$k_{DTTdeg}$ -0.6804605757196497
```

### 3. Bayesian Inferencing¶

#### Experimental Data¶

##### Upload data used for inferencing pipeline¶

- mRNA added was 0.22 μM

In [10]:

```
# Getting experimental data
folder= '/MGapt_deGFP_Exp_PURE_Data/Exp_mRNA_concs/'
filename = 'Data_0.22.csv'
mGapt_Data =  pd.read_csv(directory+ folder+ filename, delimiter = '\,', names = ['time','mRNA0', 'mRNA1','mRNA2',], engine='python',skiprows = 1)
mGapt_Data=mGapt_Data[0:144].astype(float)

filename = 'Data_0.csv'
mGapt_Data0 =  pd.read_csv(directory+ folder+  filename, delimiter = '\,', names = ['time','mRNA0'], engine='python',skiprows = 1)
mGapt_Data0=mGapt_Data0[0:144].astype(float)

#Calculating the measured MGapt
total_MGapt=pd.DataFrame(columns=['mRNA0', 'mRNA1','mRNA2'])
for name in total_MGapt.columns:
    total_MGapt[name]=((mGapt_Data[name]-mGapt_Data0['mRNA0'])-12.46)/21.39/1000
total_MGapt['time']=mGapt_Data['time']  

# Truncating data for less than 8 hours 
truncated_data = total_MGapt[total_MGapt['time']  <= 8*3600]
truncated_time = np.array(truncated_data['time'])
```

##### Getting bounded and bright states¶

In [11]:

```
# Calculating minimum fluorescence value
min_value=truncated_data[['mRNA0', 'mRNA1','mRNA2',]].T.mean().min()
altered_fluor=min_value/truncated_data.tail(1)[['mRNA0', 'mRNA1','mRNA2',]].T.mean()
altered_fluor=np.round(altered_fluor,2)

# #Defining the data for inference
df_MGapt=pd.DataFrame(columns=['mRNA0', 'mRNA1','mRNA2'])
for name in ['mRNA0', 'mRNA1','mRNA2',]:
    df_MGapt[name]=(truncated_data[name]*20-3*.22)/17
    
# Calculating percent at MGapt state
percentMGapt_t0=(np.array(df_MGapt.head(1))/.22).mean()
percentMGapt_t0=np.round(percentMGapt_t0,2)

print(f"Relative fluorescence of MGapt_altered= {float(altered_fluor)}")
print(f"Portion in of MGapt state= {float(percentMGapt_t0)}")
```

```
Relative fluorescence of MGapt_altered= 0.15
Portion in of MGapt state= 0.45
```

#### Original model¶

In [12]:

```
# Setup model
mRNA=.22 #total mRNA added by user or in reaction
initial_con={'MGapt':(mRNA_total*percentMGapt_t0), 'MGapt_altered':(mRNA_total*(1-percentMGapt_t0)),'DTT':(1000),'MGdye':(10-mRNA_total)} #in uM

modelname=filename="Model_DTTvMGapt_Final"
m = Model(sbml_filename = modelname+"_updated.xml")
m.set_species(initial_con)

# Run model to test
R0 = py_simulate_model(Model = m, timepoints = truncated_time)
R0['Measured']=R0['MGapt']+R0['MGapt_altered']*.15

#Setup plot
pMGapt=create_custom_plot('MGapt measured by Biotek', x_max=8, y_max=0.25)
pMGapt.line(truncated_time/3600, R0['Measured'], color='magenta', line_width=2,line_alpha=.4,legend_label ='Original')  
for col in ['mRNA0', 'mRNA1','mRNA2',]:
    pMGapt.scatter(truncated_time/3600, truncated_data[col], color= colors[col], radius=0.05, fill_alpha=0.2,)
pMGapt.legend.location="bottom_right"
pMGapt.legend.click_policy="hide"
pMGapt.legend.border_line_color = None
bokeh.io.show(pMGapt)
```

#### Run inference¶

In [13]:

```
#Experimental data for MGapt*
data_fit=df_MGapt.copy()
data_fit['time']=truncated_data['time']

#Setup inference
num_trajectories = 3
exp_data = []
for i in range(num_trajectories):
    df = pd.DataFrame()
    df['MGapt']=data_fit['mRNA'+str(i)]#make sure the items of measurement is the same as fit
    df["timepoints"] = data_fit["time"]
    exp_data.append(df) 

# Choose the "true" parameters.
k_MGapt  = 1.0e-3     # (k_forward) MG-dye binding to mRNA
kb_DTT   = 1.935e-06  # (k_forward1) binding of DTT to MGapt
# ku_DTT   = 7.74e-06   # (k_reverse)  unbinding of DTT to MGapt
ku_mgapt = 0.0774     # (k_forward3) unbinding of DTT from altered Malachite green aptamer
kb_mgapt = 0.07740774 # (k_reverse4) rebinding of DTT to make the altered Malachite green aptamer
# k_RNAdeg = 25.0e-4    # (k_forward5) degradation rate of unfolded mRNA
k_DTTdeg = 3.2e-4     # (k_forward6)  degradation rate of DTT

prior = {'k_forward' : ['gaussian', k_MGapt,2.4e-4 , 'positive'],
         'k_forward1' : ['gaussian',kb_DTT, 5e-7, 'positive'],
         # 'k_reverse' : ['gaussian', ku_DTT, 2e-6, 'positive'],
         'k_forward3' : ['gaussian', ku_mgapt, 0.02, 'positive'],
         'k_reverse4' : ['gaussian',kb_mgapt , 0.02, 'positive'],
         # 'k_forward5' : ['gaussian',k_RNAdeg, 20.0e-4, 'positive'],
         'k_forward6' : ['gaussian',k_DTTdeg, 5e-05, 'positive'],
         }

sampler, pid = py_inference(Model = m, exp_data = exp_data, measurements = ['MGapt'],
                            time_column = ['timepoints'], 
                            params_to_estimate = ['k_forward','k_forward1',
                                                  'k_forward3','k_reverse4','k_forward6',],
                            nwalkers = 20, nsteps = 5000, prior = prior, init_seed = 'prior', 
                            sim_type = 'deterministic', plot_show = True)
```

```
creating an ensemble sampler with multiprocessing pool= None
```

```
100%|██████████████████████████████████████████████████████████████████████████████| 5000/5000 [06:51<00:00, 12.15it/s]
```

```
Results written tomcmc_results.csv and mcmc_results.txt
Successfully completed MCMC parameter identification procedure.Check the MCMC diagnostics to evaluate convergence.
Parameter posterior distribution convergence plots:
```

In [14]:

```
# Custom plots using mcmc_results.csv
labels = ['$k_{MGapt}$','$kb_{DTT}$',
          '$ku_{mgapt}$','$kb_{mgapt}$','$k_{DTTdeg}$',]
samples_mcmc = pd.read_csv('mcmc_results.csv', names = labels,
                           engine = 'python')

corner.corner(samples_mcmc[0:], labels = labels, levels=(0.75,))
plt.savefig("Figures_DTTvMGapt/corner_plot_DTTvMGapt.svg")
```

#### Sampling over distribution¶

In [16]:

```
#Upload model
M_fit = Model(sbml_filename =  modelname+"_updated.xml")
initial_con={'MGapt':(mRNA_total*percentMGapt_t0), 'MGapt_altered':(mRNA_total*(1-percentMGapt_t0)),'DTT':(1000),'MGdye':(10-mRNA_total)} #in uM
M_fit.set_species(initial_con)

#Make plot for simulations
pfit=create_custom_plot('c', x_max=8, y_max=0.25)

#Sample over distribution
samples_discard = samples_mcmc.tail(500) #burning samples; take last 500
inds = np.random.randint(len(samples_discard), size=5)

for ind in inds:
    sample = samples_discard.iloc[ind]
    for pi, pi_val in zip(['k_forward','k_forward1',
                           'k_forward3','k_reverse4','k_forward6',], sample):
        M_fit.set_parameter(pi, pi_val)
    #Run simulation with new paramters
    fit_results = py_simulate_model(Model = M_fit, timepoints = truncated_time)   
    fit_results['Measured']=fit_results['MGapt']+fit_results['MGapt_altered']*.15
    pfit.line(truncated_time/3600, fit_results['Measured'], color='magenta', line_width=2,line_alpha=.4)    
    
for col in ['mRNA0', 'mRNA1','mRNA2',]:
    pfit.scatter(truncated_time/3600, truncated_data[col], color= colors[col], radius=0.05, fill_alpha=0.2,)
#Save of SVG
pfit.output_backend = "svg"
export_svgs(pfit, filename = 'Figures_DTTvMGapt/Fig5_pSample.svg',width=500, height=400)

bokeh.io.show(pfit)
```

#### Updated model with inferenced paramaters¶

In [17]:

```
#Upload model
M_fit = Model(sbml_filename =  modelname+"_updated.xml")
initial_con={'MGapt':(mRNA_total*percentMGapt_t0), 'MGapt_altered':(mRNA_total*(1-percentMGapt_t0)),'DTT':(1000),'MGdye':(10-mRNA_total)} #in uM
M_fit.set_species(initial_con)

#Sample over distribution
samples_mean = samples_discard.mean() #taking mean of the samples
print(samples_mean)
for pi, pi_val in zip(['k_forward','k_forward1',
                       'k_forward3','k_reverse4','k_forward6',], sample):
    M_fit.set_parameter(pi, pi_val)

#Run simulation with mean of paramters
MGapt_results = py_simulate_model(Model = M_fit, timepoints = truncated_time)   
MGapt_results['Measured']=MGapt_results['MGapt']+MGapt_results['MGapt_altered']*.15
pMGapt.line(truncated_time/3600, MGapt_results['Measured'], color='red', line_width=2,line_alpha=.4,legend_label ='Fitted')    

for col in ['mRNA0', 'mRNA1','mRNA2',]:
    pMGapt.scatter(truncated_time/3600, truncated_data[col], color= colors[col], radius=0.05, fill_alpha=0.2)
pMGapt.legend.click_policy="hide"
pMGapt.legend.location="top_left"
bokeh.io.show(pMGapt)
```

```
$k_{MGapt}$     0.001024
$kb_{DTT}$      0.000002
$ku_{mgapt}$    0.079013
$kb_{mgapt}$    0.083015
$k_{DTTdeg}$    0.000311
dtype: float64
```

### Computing environment¶

In [18]:

```
%load_ext watermark
%watermark -v -p bioscrape,bokeh,panel,jupyterlab,biocrnpyler
```

```
Python implementation: CPython
Python version       : 3.8.17
IPython version      : 8.12.2

bioscrape  : 1.2.1
bokeh      : 2.4.0
panel      : 0.13.1
jupyterlab : 3.6.5
biocrnpyler: 1.1.1
```
