## Supplementary Data Files for "Impact of Chemical Dynamics of Commercial PURE Systems on Malachite Green Aptamer Fluorescence": Model_DTTvMGapt_Sensivitiy_Inference_Plots.html

March 26, 2024

[1] Division of Engineering and Applied Science, California Institute of Technology, Pasadena, CA \

In this notebook, we plot the sensitivity and Bayesian inferencing results.

### Get data¶

In [5]:

```
# Getting experimental data
folder= '/MGapt_deGFP_Exp_PURE_Data/Exp_mRNA_concs/'
filename = 'Data_0.22.csv'
mGapt_Data =  pd.read_csv(directory+ folder+ filename, delimiter = '\,', names = ['time','mRNA0', 'mRNA1','mRNA2',], engine='python',skiprows = 1)
mGapt_Data=mGapt_Data[0:144].astype(float)

filename = 'Data_0.csv'
mGapt_Data0 =  pd.read_csv(directory+ folder+  filename, delimiter = '\,', names = ['time','mRNA0'], engine='python',skiprows = 1)
mGapt_Data0=mGapt_Data0[0:144].astype(float)

#Calculating the measured MGapt
total_MGapt=pd.DataFrame(columns=['mRNA0', 'mRNA1','mRNA2'])
for name in total_MGapt.columns:
    total_MGapt[name]=((mGapt_Data[name]-mGapt_Data0['mRNA0'])-12.46)/21.39/1000
total_MGapt['time']=mGapt_Data['time']  

# Truncating data for less than 8 hours 
truncated_data = total_MGapt[total_MGapt['time']  <= 8*3600]
truncated_time = np.array(truncated_data['time'])

#Defining values from analysis
mRNA_total=.22
percentMGapt_t0=0.45

# Model
modelname=filename="Model_DTTvMGapt_Final"
#IC for Transcription only reactions
initial_con={'MGapt':(mRNA_total*.5), 'MGapt_altered':(mRNA_total*.5),'DTT':(1000),'MGdye':(10-mRNA_total)} #in uM
```

### Sensitivity Analysis¶

In [6]:

```
#Get model
m = Model(sbml_filename = filename+"_updated.xml")
m.set_species(initial_con)

#Run sensitivity analysis
timepoints = np.linspace(0, 8*3600, 800)
ssm = py_sensitivity_analysis(model = m, timepoints = timepoints, normalize = True)
SSM=np.around(ssm[:,:,m.get_species2index()["MGapt"]].T, decimals=3, out=None)
```

```
C:\Users\zoila\anaconda3\envs\python38\lib\site-packages\bioscrape\analysis.py:244: RuntimeWarning: invalid value encountered in divide
  SSM_normalized[:,j,i] = np.divide(SSM[:,j,i]*params_values[j], solutions[:,i])
```

In [7]:

```
#Plot 
figsize = (15,7)
fig, ax = plt.subplots(figsize=figsize)
index = 1
params_names_latex = ['$k_{MGapt}$','$kb_{DTT}$','$ku_{DTT}$',
                      '$ku_{MGapt}$','$kb_{MGgapt}$','$k_{RNAdeg}$','$k_{DTTdeg}$',]

# Round the timepoints
rounded_timepoints = np.round(np.linspace(timepoints[0] / 3600, timepoints[-1] / 3600, len(timepoints),
                                           endpoint=True, dtype='float'), 1)

sn_ax = sn.heatmap(SSM, ax = ax, annot=False, cmap = 'YlGnBu',
                   xticklabels = rounded_timepoints, center=0,)

sn_ax.figure.axes[-1].xaxis.label.set_size(18)
ax = fig.axes
_ = plt.xlabel('Time (hours)', fontsize = 18)
_ = plt.ylabel('Parameters', fontsize = 18)
_ = ax[0].tick_params(axis='x', which='major', labelsize=16, bottom = False)
_ = ax[0].tick_params(axis='y', which='major', labelsize=16, left = False)
_ = ax[0].set_yticklabels(params_names_latex, rotation=0)
every_nth = 50
for n, label in enumerate(ax[0].xaxis.get_ticklabels()):
    if n == len(timepoints)-1:
        continue
    if n % every_nth != 0:
        label.set_visible(False)
_ = ax[1].tick_params(axis = 'x', labelsize = 18)
plt.savefig('Figures_DTTvMGapt/MGaptvDTT-sensitivity.svg', format='svg')
plt.show()
```

In [8]:

```
for x in range(len(params_names_latex)):
    print(params_names_latex[x], SSM[x,1:400].mean())
```

```
$k_{MGapt}$ 0.9947819548872181
$kb_{DTT}$ -0.6000200501253133
$ku_{DTT}$ 0.4551228070175438
$ku_{MGapt}$ 0.9306817042606516
$kb_{MGgapt}$ -0.9301904761904761
$k_{RNAdeg}$ -0.060438596491228055
$k_{DTTdeg}$ 1.0681553884711779
```

### Bayesian Inferencing¶

#### Plot inference results¶

In [12]:

```
# Custom plots using mcmc_results.csv
labels = ['$k_{MGapt}$','$kb_{DTT}$',
          '$ku_{MGapt}$','$kb_{MGapt}$','$k_{DTTdeg}$',]
samples_mcmc = pd.read_csv('mcmc_results_Final.csv', names = labels,
                           engine = 'python')

fig=corner.corner(samples_mcmc[0:], labels = labels, levels=(0.75,),
             label_kwargs={"fontsize": 16},)
axes = fig.get_axes()
for ax in axes:
    ax.tick_params(axis='both', which='major', labelsize=12)
# fig.savefig("Figures_DTTvMGapt/corner_plot_DTTvMGapt.svg")
```

```
WARNING:root:Pandas support in corner is deprecated; use ArviZ directly
```

#### Sampling over distribution¶

In [10]:

```
#Upload model
M_fit = Model(sbml_filename =  modelname+"_updated.xml")
initial_con={'MGapt':(mRNA_total*percentMGapt_t0), 'MGapt_altered':(mRNA_total*(1-percentMGapt_t0)),'DTT':(1000),'MGdye':(10-mRNA_total)} #in uM
M_fit.set_species(initial_con)

#Make plot for simulations
pfit=create_custom_plot('', x_max=8, y_max=0.25)

#Sample over distribution
samples_discard = samples_mcmc.tail(500) #burning samples; take last 500
inds = np.random.randint(len(samples_discard), size=5)

for ind in inds:
    sample = samples_discard.iloc[ind]
    for pi, pi_val in zip(['k_forward','k_forward1',
                           'k_forward3','k_reverse4','k_forward6',], sample):
        M_fit.set_parameter(pi, pi_val)
    #Run simulation with new paramters
    fit_results = py_simulate_model(Model = M_fit, timepoints = truncated_time)   
    fit_results['Measured']=fit_results['MGapt']+fit_results['MGapt_altered']*.15
    pfit.line(truncated_time/3600, fit_results['Measured'], color='magenta', line_width=2,line_alpha=.4)    
    
for col in ['mRNA0', 'mRNA1','mRNA2',]:
    pfit.scatter(truncated_time/3600, truncated_data[col], color= colors[col], radius=0.05, fill_alpha=0.2,)
#Save of SVG
pfit.output_backend = "svg"
# export_svgs(pfit, filename = 'Figures_DTTvMGapt/Fig5_pSample.svg',width=500, height=400)

bokeh.io.show(pfit)
```

### Computing environment¶

In [11]:

```
%load_ext watermark
%watermark -v -p bioscrape,bokeh,panel,jupyterlab,biocrnpyler
```

```
Python implementation: CPython
Python version       : 3.8.17
IPython version      : 8.12.2

bioscrape  : 1.2.1
bokeh      : 2.4.0
panel      : 0.13.1
jupyterlab : 3.6.5
biocrnpyler: 1.1.1
```
