## Supplementary Data Files for "Impact of Chemical Dynamics of Commercial PURE Systems on Malachite Green Aptamer Fluorescence": 2023.09.09_SNARF-5F_PURExpress.ipynb

```
# %% [code]
#workhorses
import numpy as np
import pandas as pd
import math
import seaborn as sns; sns.set()
import matplotlib.pyplot as plt
import random 
from bokeh.models import Label
from scipy import stats

import bokeh
import matplotlib.pyplot as plt
# bokeh.io.output_notebook()

import holoviews as hv
hv.extension('bokeh')
# bebi103.hv.set_defaults()
from sklearn.metrics import r2_score
# from bokeh.io import gridplot, output_file, show
from bokeh.io import export_png
from bokeh.models import Title
from bokeh.plotting import gridplot,figure, output_file, show
from bokeh.models.glyphs import Text

#for custom colormaps
from matplotlib.colors import LinearSegmentedColormap

#Get directory
import os
directory = os.getcwd()

# %% [markdown]
## Bokeh

# %% [code]
%matplotlib inline
import bokeh.io
import bokeh.plotting
bokeh.io.output_notebook()
from bokeh.themes import Theme

# Modules needed from Bokeh.
from bokeh.io import output_file, show
from bokeh.plotting import figure
from bokeh.models import LinearAxis, Range1d
from bokeh.palettes import Magma, Inferno, Plasma, Viridis, Cividis

colors = bokeh.palettes.Colorblind[8]
colors2= bokeh.palettes.Magma[11]

try:
    import dnaplotlib as dpl
    dpl_enabled = True
except (ModuleNotFoundError,ImportError) as e:
    dpl_enabled = False
    
theme = Theme(json={'attrs': {
# apply defaults to Figure properties
'Figure': {
    'toolbar_location': 'right',
    'outline_line_color': None,
    'min_border_right': 10,
#     'sizing_mode': 'stretch_width',
    'height':600,
    'width':800,
},'Grid': {
    'grid_line_color': None,
},
'Title': {
    'text_font_size': '20pt',
    'align': 'center'
},
    
    
# apply defaults to Axis properties
'Axis': {
#     'minor_tick_out': None,
#     'minor_tick_in': None,
    'major_label_text_font_size': '15pt',
    'axis_label_text_font_size': '15pt',
#     'axis_label_text_font': 'Work Sans',
    'axis_label_text_font_style': 'normal',
    'axis_label_standoff':15
},


# apply defaults to Legend properties
'Legend': {
    'background_fill_alpha': 0.8,
    'location': 'top_right',
    "label_text_font_size": '15pt'
}}})

bokeh.io.curdoc().theme = theme
from bokeh.io import export_png

# %% [code]
#plotting things

#%matplotlib qt5 -- I don't know what this is
import matplotlib.pyplot as plt
import matplotlib as mpl
import seaborn as sns

from cycler import cycler


#All of Anandh's customized seaborn/matplotlib settings

sns.set_context("talk", font_scale=1.5, rc={"lines.linewidth": 1.5})
sns.set_style("ticks")
sns.set_style({"xtick.direction": "in","ytick.direction": "in"})

#%config InlineBackend.figure_f.ormats=['svg']

mpl.rc('axes', prop_cycle=(cycler('color', ['r', 'k', 'b','g','y','m','c']) ))

mpl.rcParams['pdf.fonttype'] = 42
mpl.rcParams['ps.fonttype'] = 42

#mpl.rc('text', usetex=False)
#mpl.rc('text.latex', preamble=r'\usepackage{helvet}
#\renewcommand\familydefault{\sfdefault}\usepackage{sansmath}\sansmath')

    #If you want to use a different font
# mpl.rc('font',**{'family':'sans-serif','sans-serif':['Helvetica'], 
#                  'serif': ['Helvetica']})

tw = 1.5
sns.set_style({"xtick.major.size": 3, "ytick.major.size": 3,
               "xtick.minor.size": 2, "ytick.minor.size": 2,
               'axes.labelsize': 16, 'axes.titlesize': 16,
               'xtick.major.width': tw, 'xtick.minor.width': tw,
               'ytick.major.width': tw, 'ytick.minor.width': tw})

mpl.rc('xtick', labelsize=14) 
mpl.rc('ytick', labelsize=14)
mpl.rc('axes', linewidth=1.5)
mpl.rc('legend', fontsize=14)

mpl.rc('figure', figsize=(11,9))


# %% [markdown]
# Munge Data

# %% [markdown]
## Definitions Needed

# %% [markdown]
### Concat multiple plates over time

# %% [code]
def fix_columns (dataframe):
    
    renames = {curr_name : dataframe.loc[min(dataframe.index), curr_name] for curr_name in dataframe.columns}
    
    dataframe = dataframe.rename(columns = renames)
    
    dataframe = dataframe.drop(min(dataframe.index))
    
    return dataframe

# %% [code]
def get_plate_dfs (multiplate_df):
    
    #gets the indices where the row says "time" which is the header row for that plate's table
    times = multiplate_df.loc[multiplate_df['Time'] == 'Time'].index
    
    #since the first one has it's header as the column titles already, add this index to capture the first one
    timesn = [0] + list(times) + [max(multiplate_df.index) + 1]
    
    #get number of indices to add to the "Times" header row to get this plate's table
    timediffs = np.diff(np.array(timesn))
    
    #get the start and end indices for each table you want
    start_end_idxs = [(i, i+j) for i,j in zip(timesn, timediffs)]
    
    #select the data from the big composite set
    datas = [multiplate_df.iloc[x[0]:x[1]] for x in start_end_idxs]
    
    #the first set has proper column labels, the others have the column labels as the first row
    #replace the existing columns with the first row entries to get proper column labels
    alls = [datas[0]] + [fix_columns(data) for data in datas[1:]]
    
    #put them all together in order to get the final df
    #concat joins them on matching columns, so order of columns doesn't matter, just that they are present in all dfs
    output = pd.concat(alls).reset_index(drop=True)
    
    return output

# %% [markdown]
### Munging

# %% [code]
def get_sec (datetimeTime_obj):
    
    return (datetimeTime_obj.hour * 60 + datetimeTime_obj.minute) * 60 + datetimeTime_obj.second

# %% [code]
def replace_time (data_sheet):
    
    df = data_sheet.copy()
    
    df['Time'] = df['Time'].apply(get_sec).divide(3600)
    
    return df

# %% [code]
def replace_time_sequential (data_sheet):
    
    df = data_sheet.copy()
    
    df['Time'] = df['Time'].apply(get_sec).divide(3600)
    
    first = df.loc[0, 'Time']
    second = df.loc[1, 'Time']
    
    diff = second - first
    
    too_long = np.arange(first, 1000, diff)
    
    proper_length = too_long[:(len(df['Time']))]
    
    df['Time'] = proper_length
    
    return df

# %% [code]
def replace_temp (data_sheet):
    
    df = data_sheet.copy()
    
    c = ['Temp C' if '°' in x else x for x in df.columns]
    
    df.columns = c
    
    return df

# %% [code]
def add_id_info (data_sheet_melted, df_id):
    
    #make a copy of the dataframe so you can return the new one and set whatever name you want
    df = data_sheet_melted.copy()

    #initialize the columns to be used to store ID info
    all_id_columns = [col for col in df_id.columns if col not in 'well']

    for c in all_id_columns:
        df[c] = np.nan

    #get the list of all the wells you want to annotate
    wells = np.unique(df_id['well'])

    #loop over these wells and add the information to the initialized columns
    for w in wells:
        #get the ID information for this well
        append_this = df_id.loc[df_id['well'] == w, :]

        #loop over each column containing a unique piece of ID info
        for c in all_id_columns:

            #set the value for that column for that well
            df.loc[df['well'] == w, c] = append_this[c].values
                                                                #have to use .values otherwise index carries along and only
                                                                #sets the value for the data table index equal to the
                                                                #id table index

    return df

# %% [code]
def add_id_info (data_sheet_melted, df_id):
    
    #make a copy of the dataframe so you can return the new one and set whatever name you want
    df = data_sheet_melted.copy()

    #initialize the columns to be used to store ID info
    all_id_columns = [col for col in df_id.columns if col not in 'well']

    for c in all_id_columns:
        df[c] = np.nan

    #get the list of all the wells you want to annotate
    wells = np.unique(df_id['well'])

    #loop over these wells and add the information to the initialized columns
    for w in wells:
        #get the ID information for this well
        append_this = df_id.loc[df_id['well'] == w, :]

        #loop over each column containing a unique piece of ID info
        for c in all_id_columns:

            #set the value for that column for that well
            df.loc[df['well'] == w, c] = (append_this[c].reset_index(drop=True))[0]
                                                                #have to use .values otherwise index carries along and only
                                                                #sets the value for the data table index equal to the
                                                                #id table index
    return df

# %% [markdown]
## get data

# %% [code]
filename = '/2023.09.09_SNARF-5F_PURExpress.xlsx'
data_dict = pd.read_excel(directory + filename, sheet_name=None, engine='openpyxl')
sheets=data_dict.keys()

# %% [code]
sheets_to_tidy= [x for x in data_dict.keys() if '_data' in x]
sheets_to_tidy

# %% [markdown]
### Assign well IDs

# %% [code]
ids = data_dict['IDs']

#get the data you want
for s in sheets_to_tidy:
    s_raw=data_dict[s]
    s_raw=replace_time(s_raw)
    name= s
    #independently melt the data so you can check and control the ID vars and value vars since that might differ per expt
    df = pd.melt(s_raw, id_vars=['Time','Temp C'], var_name=['well'])
    df = add_id_info(df, ids)
    #Write the data to a new file
    df.to_csv(s + '_tidy.csv', index=False) 

# %% [markdown]
# Computing environment

# %% [code]
%load_ext watermark
%watermark -v -p bioscrape,bokeh,panel,jupyterlab,biocrnpyler
```
